# Site-Specific Labelling of Multidomain Proteins by Amber Codon Suppression

**DOI:** 10.1101/282525

**Authors:** Christina S. Heil, Alexander Rittner, Bjarne Goebel, Daniel Beyer, Martin Grininger

## Abstract

Amber codon suppression is a powerful tool to site-specifically modify proteins to generate novel biophysical probes. Yet, its application on large and complex multidomain proteins is challenging, leading to difficulties during structural and conformational characterization using spectroscopic methods. The animal fatty acid synthase type I is a 540 kDa homodimer displaying large conformational variability. As the key enzyme of *de novo* fatty acid synthesis, it attracts interest in the fields of obesity, diabetes and cancer treatment. Substrates and intermediates remain covalently bound to the enzyme during biosynthesis and are shuttled to all catalytic domains by the acyl carrier protein domain. Thus, conformational variability of animal FAS is an essential aspect for fatty acid biosynthesis. We investigate this multidomain protein as a model system for probing amber codon suppression by genetic encoding of non-canonical amino acids. The systematic approach relies on a microplate-based reporter assay of low complexity, that was used for quick screening of suppression conditions. Furthermore, the applicability of the reporter assay is demonstrated by successful upscaling to both full-length constructs and increased expression scale. The obtained fluorescent probes of murine FAS type I could be subjected readily to a conformational analysis using single-molecule fluorescence resonance energy transfer.

## Introduction

Fatty acid synthases type I (FASs) are large and complex multidomain enzymes that are responsible for cytosolic *de novo* fatty acid synthesis^1,2^. Evolutionarily related to FAS are polyketide synthases type I (PKSs), that synthesize polyketides, which account for one of the largest classes of natural products^3,4^. In FASs and PKSs, multiple catalytic sites interact successively to stepwise assemble fatty acids or the complex and chemically diverse polyketides (Fig. 1A)^5^. A common feature of these enzymes is, that the substrates remain covalently bound to the acyl carrier protein (ACP) domain during catalysis. The ACP domain interacts with all catalytic domains, which requires large positional variability within the FAS and PKS systems^6–9^.

**Figure 1:**
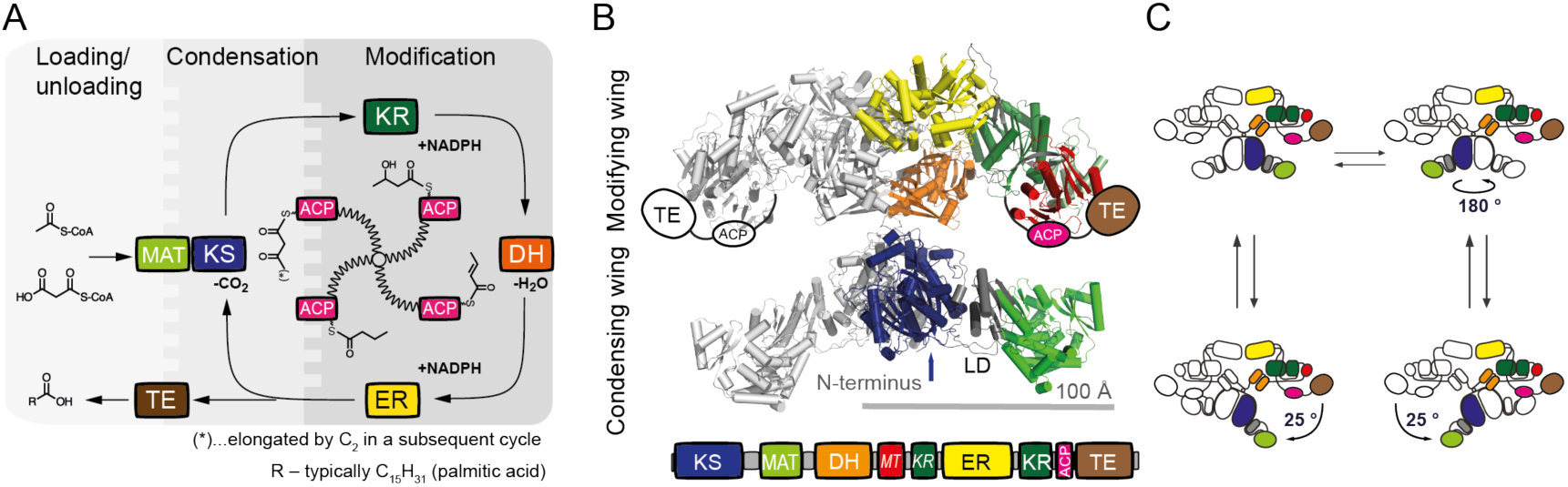
Overview of animal fatty acid synthesis. A) Fatty acid synthesis as occurring in animals. The fatty acid, typically palmitic acid, is produced from the substrates acetyl-CoA, malonyl-CoA and NADPH. The acetyl moiety is sequentially elongated and modified by several domains until a certain chain length (C_16_) is reached and the final product is released from the enzyme as a free fatty acid. During the whole process all intermediates remain covalently attached to the enzyme, mainly to the ACP domain, which requires a high conformational freedom of FAS to facilitate productive interactions between the ACP domain and all catalytically active sites. Domain nomenclature: KS (ketoacyl synthase), KR (ketoacyl reductase), DH (dehydratase), ER (enoyl reductase), ACP (acyl carrier protein), TE (thioesterase), MAT (malonyl/acetyltransferase). B) Cartoon depiction of the dimeric “X”- shaped structure of porcine FAS^6^. α-Helices are shown as cylinders. One half of the dimer is coloured according to the attached domain overview. Owing to their high positional variability, ACP and TE could not be traced in electron density, but are schematically drawn for clarity. *KR* and *MT* (methyltransferase) refer to non-catalytic folds, which have structural tasks and may confine the ACP during substrate shuttling. C) Conformational dynamics of animal FAS. Swinging and swivelling motions around the flexible hinge region have been observed by single particle EM and high-speed atomic force microscopy^8,15^. Full rotation of the condensing wing by 180° was further confirmed by mutagenesis studies^50^.

While the overall architecture of type I PKSs has not yet been firmly elucidated^10,11^, high resolution data of FASs in different structural arrangements are available^6,12^. As observed in 3.2 Å model X-ray crystal structure on FAS from pig, animal FAS assembles into an intertwined dimer of approximately 540 kDa, adopting an “X”-shaped conformation (Fig. 1B). Although the ACP and TE domains could not be traced in the electron density, it becomes apparent from the model, that a positionally variable ACP alone is not able to reach every catalytic centre. This paradox was already described by Hammes *et al*. in an early fluorescence resonance energy transfer (FRET) study on chicken liver FAS^13,14^. More detailed insights into the conformational versatility of animal FAS were finally given by a recent negative stain electron microscopy (EM) study on rat FAS, high-speed atomic force microscopy on insect FAS and by computational modelling with porcine FAS data^7,8,15^. These studies revealed large conformational changes within the enzyme, with complete relative rotational and swinging freedom between the condensing and processing wing (Fig. 1C).

To study the conformational dynamics of animal FAS and related PKSs in real-time, we seek to establish spectroscopic methods at the single-molecule level, as they can deliver a continuous spectrum of stochastic conformational motions in proteins^16,17^. An integral aspect of spectroscopic methods is the modification of proteins with labels. Conventional techniques, such as labelling naturally occurring or mutationally introduced cysteines via maleimide chemistry^18^, are not applicable for animal FAS, since the large complex features many native cysteines, including active site cysteines. Our method of choice was therefore the genetic encoding of non-canonical amino acids (ncAAs) through the amber codon suppression technology^19,20^. Such ncAAs carry orthogonal functional groups, which can be used to site-specifically attach spectroscopic labels by post-translational modification.

To the best of our knowledge, the introduction of ncAAs and the subsequent bioconjugation with a fluorophore have neither been reported for megasynthases nor for any other multidomain protein of such sizes to date. We therefore established a systematic approach, in which the screening of amber codon suppression systems, ncAA insertion sites and fluorophore click protocols can be performed with an authentic system of low complexity, that promises a high success rate in upscaling for the production of the full-length protein. Specifically, we set up a microplate-based reporter assay and upscaling protocols, which achieved the production of ncAA- modified murine FAS (mFAS), further successfully labelled with fluorophores. The reporter assay, performed on an ACP-GFP fusion protein, is a suited platform for the screening of ncAA incorporation by the read-out of the fluorescence of the fused GFP domain^21^. Successful upscaling demonstrates the reliability and applicability of reporter assay data to larger protein constructs in increased expression scale.

## RESULTS

### Screening of the amber codon suppression toolbox

Out of a growing repertoire of ncAAs, that are introduced by amber codon suppression, we limited our screening to eight different ncAAs with functional groups, that can be used for bioconjugation with click chemistry or oxime formation (Fig. 2A; for syntheses of respective ncAAs, see Supplementary Methods)^19^. Azido- and propargyl-functional groups can for example be used in copper(I)-catalysed alkyne-azide cycloadditions (CuAACs)^22^. Since copper is critical for the stability of the protein, we focused on copper-free click chemistry, like the strain-promoted alkyne-azide cycloaddition (SPAAC)^23^, with azido- and bicyclononyne (BCN)-functional groups, and the inverse electron-demand Diels-Alder cycloaddition (IEDDAC)^24^, with tetrazine- and norbonene-functional groups. Additionally, we also tested incorporation of the ncAA AcLys, as the acyl-functional group can be bioconjugated in oxime formation^25^, and acetylation of lysines naturally occurs as post-translational modification in animal FAS^26^.

**Figure 2:**
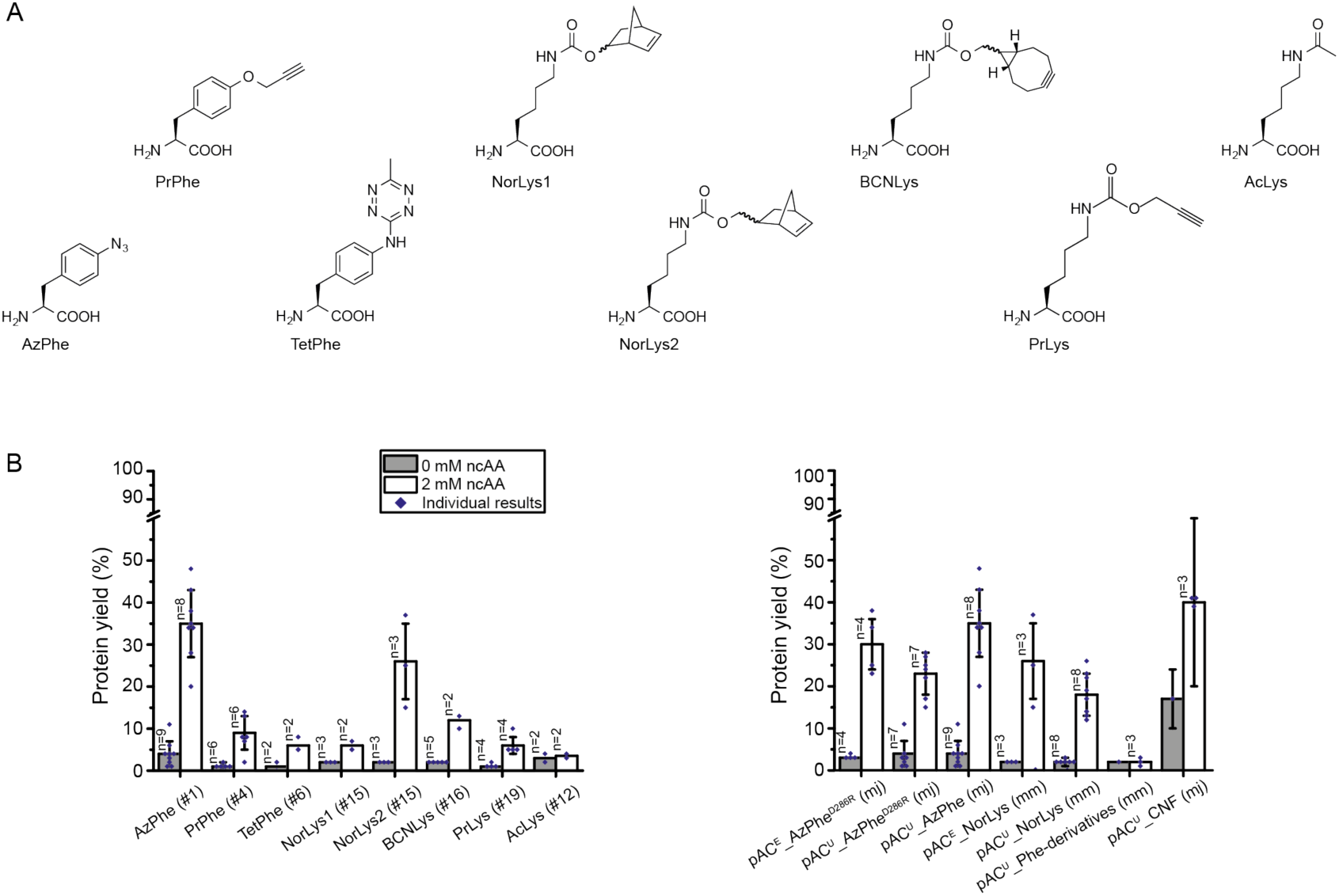
Amber codon suppression at site Leu54 in the ACP-GFP fusion construct screened in reporter assay. A) Overview of ncAAs used in this study. B) Best expression efficiency of different ncAAs (left panel) and comparison of some representatives of the screening (right panel). Respective plasmids used for incorporation of ncAAs are indicated by plasmid number (#; listed in Supplementary Table S2). A compilation of all results from the reporter assay can be found in Supplementary Fig. S2. Expression efficiency is read out by GFP fluorescence of 2 mL *E. coli* cell cultures and compared to wild type reference (taken as 100%). For incorporation, 2 mM ncAAs were supplemented to the medium. Cultures lacking ncAAs were taken as negative control to determine background signal. Dots refer to values of the biological samples. The averages of biological samples are plotted together with standard deviations. Technical errors were below 10%.

Further, we compared two common suppressor vectors pUltra and pEVOL (see Supplementary Fig. S1), published by Schultz and coworkers^27,28^, and several evolved aminoacyl-tRNA synthetase (aaRS)/tRNA pairs from *Methanococcus jannaschii, Methanosarcina mazei* and *Methanosarcina barkeri* for their performance. The cloning procedure of suppressor vectors pAC^U^ and pAC^E^ (based on original pUltra and pEVOL, respectively) is described in detail in the Supplementary Methods. Supplementary Table S1 lists all primers used for cloning, and Supplementary Table S2 summarizes all suppressor plasmids and evolved aaRS of this study.

To identify the optimal pair of suppression system and ncAA, we established a reporter assay. The screening was performed on a fusion construct of ACP from mFAS with GFP (ACP-GFP), placing the amber mutation in a disordered and non-conserved loop region at the Leu54 site, using the homologous rat ACP structure (PDB: 2png) as template. Incorporation efficiency was read out at 2 mL scale by recording the fluorescence of *E. coli* cell cultures expressing different ACP-GFP mutants. Cultures lacking the ncAA in the medium were taken as negative controls to determine the background signal (Fig. 2B). Negative samples showed a fluorescence level of up to 4% of the wild type reference. High incorporation efficiencies were observed for AzPhe^29^ (35% ± 8%) and NorLys2^30,31^ (26% ± 9%). The ncAA BCNLys^32^ was incorporated with 12% ± 2% efficiency, all other ncAAs (PrPhe^33^, TetPhe^34^, NorLys1^24^ and PrLys^35^) showed efficiencies below 10%, and AcLys^36^ was hardly incorporated at all. Comparing the suppressor plasmids pAC^E^ and pAC^U^, the plasmid pAC^E^ seemed to be slightly more efficient in our set-up than its pAC^U^ counterpart (30% ± 6% pAC^E^_AzPhe^D286R^ vs. 23% ± 5% pAC^U^_AzPhe^D286R^ and 26% ± 9% pAC^E^_NorLys vs. 18% ± 5% pAC^U^_NorLys). The D286R mutation in the aminoacyl-tRNA synthetase of *M. jannaschii*^37^, which was reported to have a beneficial effect, did not improve incorporation efficiencies in our hands (23% ± 5% pAC^U^_AzPhe^D286R^ vs. 35% ± 8% pAC^U^_AzPhe). Comparing the two orthogonal systems, the tyrosyl-tRNA synthetase derived from *M. jannaschii* (mjTyrRS) seemed to be more efficient in our set-up than the pyrrolysyl-tRNA synthetases of *M. mazei* or *M. barkeri* (mmPylRS or mbPylRS, respectively) (e.g. 35% ± 8% pAC^U^_AzPhe (mj) vs. 18% ± 5% pAC^U^_NorLys (mm)). We also tested two less specific suppressor vectors, which incorporate multiple ncAAs. While the suppressor plasmid pAC^U^_Phe-derivatives of *M. mazei*^38^ failed to incorporate any phenylalanine derivatives, the suppressor plasmid pAC^U^_CNF from *M. jannaschii* ^39,40^ showed high incorporation efficiencies of 40% ± 11%, but suffered from relatively high fluorescence of the negative control (18% ± 7%), indicating unspecific incorporation of endogenous amino acids. A compilation of all results from the reporter assay screening can be found in Supplementary Fig. S2.

### Screening of ncAA incorporation sites

As it has been reported before that the specific position of an amber mutation has major effects on incorporation efficiencies^41^, we used the most promising systems from the initial screening to compare incorporation efficiencies at different sites (pAC^U^_AzPhe with AzPhe as the optimal result and the respective vector pAC^U^_NorLys for NorLys2). Hence, we selected six positions in the ACP fold to test the acceptance of ncAA incorporation (Ala in the linker region between the N-terminal Strep-tag and ACP-GFP, Gly01 at the N-terminus of the mouse ACP sequence, Leu54 in a disordered loop region, Gln70 and Asp71 in helix 5 and Ala79 in the linker between ACP and GFP; see Fig. 3A and 3B). AzPhe was incorporated with good efficiencies (in average 38% ± 1%) throughout all amber mutation sites, whereas the incorporation efficiencies for NorLys2 were strongly dependent on the respective position (Fig. 3C). Best incorporation efficiency for NorLys2 was gained for the amber mutation site Gly01 at the N-terminus with 39% ± 13%. The amber mutation site Leu54 in the disordered loop region of ACP showed 16% ± 6% incorporation efficiency for NorLys2, whereas amber mutation site Gln70 in the last helix of ACP showed no incorporation at all. All other amber mutation sites showed incorporation efficiencies below 10%. Higher concentrations (4 mM and 8 mM) of ncAAs in the medium seemed to slightly increase incorporation efficiency of NorLys2 and slightly decrease efficiency of AzPhe (see Supplementary Fig. S3). Therefore, we proceeded with a concentration of 2 mM ncAA.

**Figure 3:**
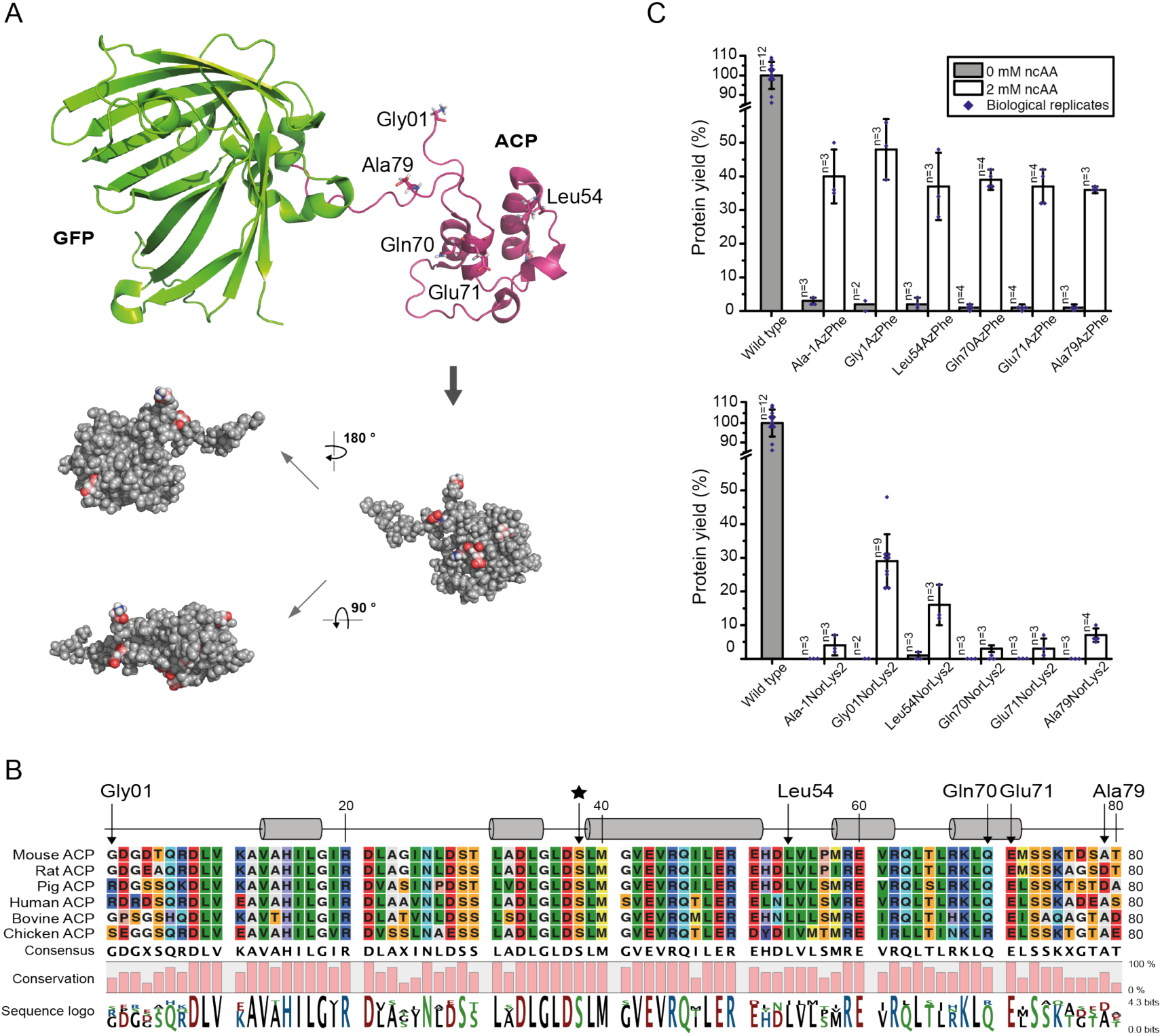
Screening of amber codon mutation sites. A) Cartoon representation of the ACP-GFP fusion construct (upper panel; pink: rat ACP (PDB: 2png) and green: eGFP (PDB: 2y0g)) used in the reporter assay. The five amber mutation sites are labelled and depicted in stick representation (Gly01, Leu54, Gln70, Glu71 and Ala79). Different orientations of the ACP domain (shown in a sphere-filling model) demonstrate the positioning of all amber mutation sites on the surface of the domain (lower panel). Amber mutation sites are coloured in red. B) Sequence alignment of six different ACP domains of animal FASs. Uniprot accession codes: mouse FAS: P19096; rat FAS: P12785; pig FAS: A5YV76; human FAS: P49327; bovine FAS: Q71SP7 and chicken FAS: P12276. The five amber mutation sites are highlighted by arrows, and a star highlights the active serine residue. Secondary structure elements received from the rat ACP model (PDB: 2png) are depicted (α-helices shown as cylinders). C) Expression efficiencies of six different AzPhe mutants (upper panel) and six different NorLys2 mutants (lower panel) in comparison to the wild type reference, read out by the GFP fluorescence of 2 mL cultures of *E. coli* cells. For incorporation, 2 mM ncAAs were supplemented to the medium. Cultures lacking ncAAs were taken as negative control to determine background signal. The averages of biological replicates are plotted together with standard deviations and the distribution of individual values is indicated as dots. Technical errors were below 10%.

### Upscaling of protein production

In a next step, it was tested whether the selected conditions from the reporter assay could be reproduced in larger expression cultures of 200 mL. Each culture was analysed in fluorescence, as was implemented in the reporter assay, and further evaluated by the yield of purified protein. Fluorescence data was collected similarly to the reporter assay, taking a 2 mL sample of the cell culture. The ncAA AzPhe was incorporated with overall good efficiency (in average 32% ± 4%), whereas large variations were observed for incorporation of NorLys2 at different amber mutation sites (Fig. 4A). Best incorporation efficiency for NorLys2 was gained for the amber mutation site Leu54 with 29% ± 3% and no incorporation was achieved at amber mutation site Gln70. This data agreed well with the results from the reporter assay, except for the incorporation of NorLys2 in Gly01 failing at larger volume, while leading to best incorporation efficiencies in the reporter assay. We observed systematic higher values for NorLys2 and slightly lower values for AzPhe by GFP-fluorescence in the larger expression culture as compared to the reporter assay.

**Figure 4:**
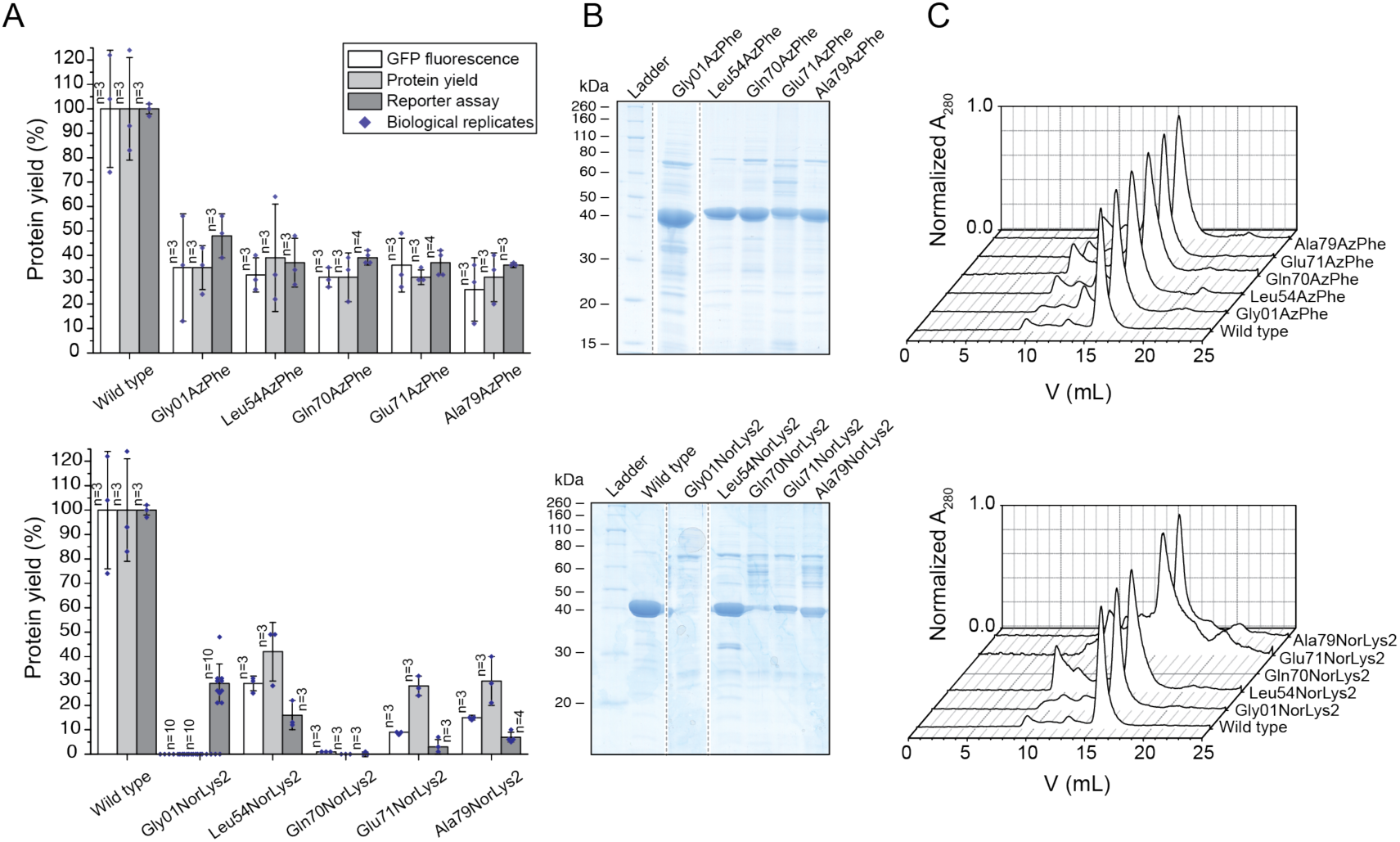
Large scale expression and purification of ACP-GFP mutants (upper panels AzPhe mutants, lower panels NorLys2 mutants). A) Comparison of the results from large scale expression cultures (protein yield was read out by GFP fluorescence of a 2 mL sample and by the yield of purified protein) with previous results from the reporter assay. Data compare expression efficiency of wild type and five different AzPhe mutants (upper panel), and expression efficiency of wild type and five different NorLys2 mutants (lower panel). All expression efficiencies are related to the wild type reference. For incorporation, 2 mM ncAA were supplemented to the medium. The averages of biological replicates are plotted together with standard deviations and the distribution of individual values is indicated as dots. Technical errors were below 10%. B) SDS-PAGE (NuPAGE Bis-Tris 4-12%) gel of ACP-GFP mutants purified by Ni-chelating chromatography. Lanes have been assembled for clarity, but scans of the original gels can be found in Supplementary Fig. S6. SDS-PAGE shows one set of purified proteins (one biological replicate). C) Preparative SEC of ACP-GFP mutants performed with a Superdex 200 Increase 10/300 GL column (the set of proteins shown in B). Peaks at an elution volume of 16 mL correspond to the ACP-GFP variants. The void volume of the column is at ca. 9 mL.

For comparing protein yields, cells received from 200 mL expression cultures were lysed and proteins were isolated by Ni-chelating chromatography. Compared to ACP-GFP at 53 ± 15 mg, the positive mutants were expressed in average with 14 ± 3 mg yield, which corresponds to about 25% of the wild type protein yield (Fig. 4B). The incorporation efficiency quantified by protein yield correlated well with the trends of the fluorescence data. AzPhe was again incorporated with overall good efficiency (in average 33% ± 4%), whereas NorLys2 performed differently throughout the amber mutation sites (Fig. 4A). The optimal site for NorLys2, Leu54 in the disordered loop region, showed 42% ± 12% incorporation efficiency, and even the amber mutation sites Glu71 and Ala79 gave up to 30% incorporation efficiency. Again, no incorporation of NorLys2 at the amber mutation site Gln70 was monitored. We note that NorLys2 mutants led to higher protein yields than expected from fluorescence intensities of cell cultures, which cannot be explained with the collected data. The quality of proteins was analysed by size exclusion chromatography (SEC) and mass spectrometry (MS). The elution profiles of the different ACP-GFP mutants matched very well with the wild type SEC spectrum (Fig. 4C). The incorporation of the ncAAs was confirmed by MS analysis (see Supplementary Data).

### Fluorescent labelling of ACP-GFP

In first experiments, we screened click kinetics for our target protein ACP-GFP (see Supplementary Fig. S4) and received a suited condition of 2 h of incubation at room temperature with 80 equiv. of fluorophore in 10 µL reaction volume for both the SPAAC (AzPhe mutant BCN-POE_3_-NH-DY649P1 conjugate) and the IEDDAC (NorLys2 mutant 6-methyl-tetrazine-ATTO-647N conjugate). For determining the degree of labelling (DOL) by in-gel fluorescence, ACP-GFP enzymatically modified by a fluorescent CoA-label by a 4’-phosphopantetheinyl transferase (Sfp)^42^ was used as reference. For better comparison, the different fluorescence intensities were corrected by the respective quantum efficiencies. Three different fluorophores were used in this experiment: DY647P1 at the CoA-label (quantum efficiency 30% according to the manufacturer), DY649P1 at the BCN-label (quantum efficiency 30% according to the manufacturer) and ATTO 647N at the tetrazine-label (quantum efficiency 65% according to the manufacturer). The sample of ACP-GFP enzymatically modified by a fluorescent CoA-label showed highest fluorescence and was assumed to be quantitatively labeled^43^, and thus set to 100% as the wild type reference. All fluorescence intensities were further correlated to the intensity of the protein bands of the Coomassie-stained gel. In average, the AzPhe mutants clicked more efficiently than the NorLys2 mutants (74% over 23%) (Fig. 5A).

**Figure 5:**
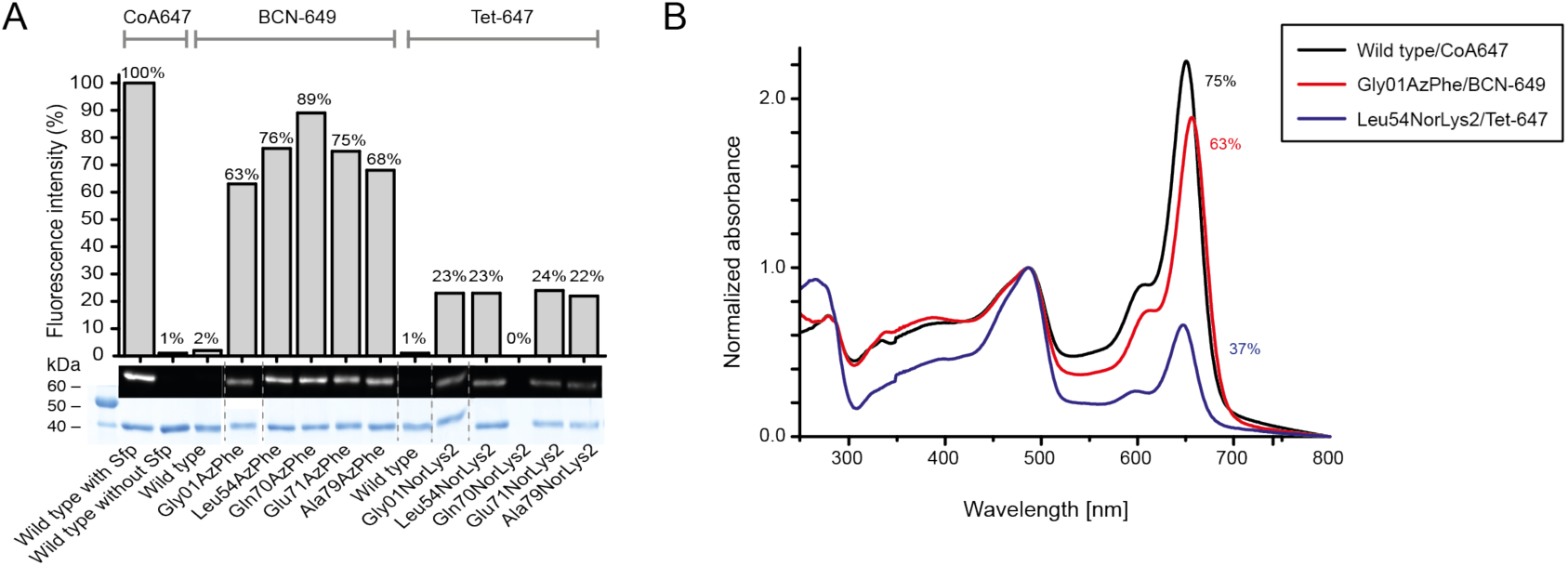
Fluorescent labelling of ACP-GFP mutants. A) DOL of ACP-GFP mutants in respect to the amber mutation site determined by relative in-gel fluorescence intensities at wavelength 650 nm. The ACP-GFP construct was enzymatically modified by a fluorescent CoA647-label with Sfp and served as the wild type reference. Hence, it was put to 100% fluorescence intensity. AzPhe mutants were labelled with 80 equiv. of BCN-POE_3_-NH- DY649P1 (BCN-649), NorLys2 mutants were labelled with 80 equiv. of 6-methyl-tetrazine-ATTO-647N (Tet-647) in 10 µL reaction volume. All fluorescence intensities were corrected by the quantum efficiency of the respective fluorophore and correlated to the protein bands of the Coomassie-stained gel (lanes have been assembled for clarity). Scans of the original gels are presented in Supplementary Fig. S7. B) DOL determined by UV-Vis spectroscopy. 25 equiv. of fluorophore were used in labelling reactions of ACP-GFP mutants in 50 µL reaction volume. Free fluorophore was removed by purification over Ni-NTA magnetic beads. UV-Vis absorbance spectra were normalized to GFP absorbance at wavelength 485 nm. DOL is read out by comparing absorbance of the fluorophore at 650 nm to absorbance of GFP at 485 nm.

The DOL was alternatively determined by spectroscopy with samples Gly01AzPhe and Leu54NorLys2, after removal of excess free fluorophore by purification over Ni-NTA magnetic beads. In a single experiment, these proteins were clicked with 25 equiv. of fluorophore in 50 µL reaction volume and the labelling efficiency was monitored with UV-Vis absorbance spectra. For the wild type reference, a DOL of 75% was determined, whereas the Gly01AzPhe mutant showed 63% DOL and the Leu54NorLys2 mutant showed only 37% DOL (Fig. 5B). We explain the difference in DOL as originating from different reaction conditions and sample preparations performed for analysis in SDS-PAGE and spectroscopy (see Fig. 5A and B). The quantum efficiency is determined for the free fluorophore and may be differently affected by the microenvironment within the native and the denatured protein. Data may also indicate that ACP was not quantitatively labelled during enzymatic modification with fluorescent CoA-label. As in-gel fluorescence is always determined relative to the wild type reference, accurate comparison of intensities between different gels is difficult, and thus we observed variations in the DOL in further labelling reactions. We note that the DOL was determined in single experiments by two different methods, without the claim for statistical representation.

### Application of selected conditions on full-length mFAS

Three amber mutations were introduced in the ACP domain of full-length mFAS. Two of those mutations were selected as promising candidate constructs at positions Gly2113 (Gly01 in ACP) and Leu2166 (Leu54 in ACP), and the other mutation as a negative candidate construct at position Gln2182 (Gln70 in ACP). The incorporation of ncAAs to yield the proteins Gly2113AzPhe, Leu2166NorLys2 and Gln2182NorLys2 was performed with the conditions identified by screening and upscaling experiments, as described above. In agreement with data of the reporter assay and ACP-GFP protein analysis, the mutants Gly2113AzPhe and Leu2166NorK2 were expressed in good yields (1.9 mg/L culture and 0.9 mg/L culture, respectively), whereas the Gln2182NorK2 mutant expressed poorly (0.03 mg/L culture, in comparison to the wild type mFAS: 2.9 mg/L culture) (see Supplementary Fig. S5). Western blot analysis with antibodies against the N-terminal Strep-tag and C-terminal His-tag was used for reading out incorporation efficiency. Double bands are observed for the mutants Gly2113AzPhe and Leu2166NorK2. This indicates mixtures of mFAS in full-length (upper red & green band) and truncated protein (lower red band) (both bands at about 260 kDa size), indicating probably insufficient suppression of the amber codon (Fig. 6A). Western Blot analysis of the negative candidate Gln2182NorK2 mutant shows only the lower molecular weight band corresponding to truncated band, and indicates the failed incorporation of the ncAA. The purified constructs of mFAS were further clicked with fluorophores, which confirmed successful incorporation of the ncAAs. In addition to specific labelling of the protein (by clicking complementary fluorophores), a small amount of unspecific binding was detected (Fig. 6B). Again, the SPAAC (Gly2113AzPhe mutant BCN-POE_3_-NH-DY649P1 conjugate) seems to be more efficient than the IEDDAC (Leu2166NorLys2 mutant 6-methyl-tetrazine-ATTO-647N conjugate) (Fig. 6B). Fluorescence detected for Gly2113AzPhe mutant BCN-POE_3_-NH-DY649P1 conjugate is higher than for the wild type CoA 647 conjugate, which may be explained with incomplete phosphopantetheinylation of the reference protein and some amount of unspecific binding of the BCN-POE_3_-NH-DY649P1 fluorophore.

**Figure 6:**
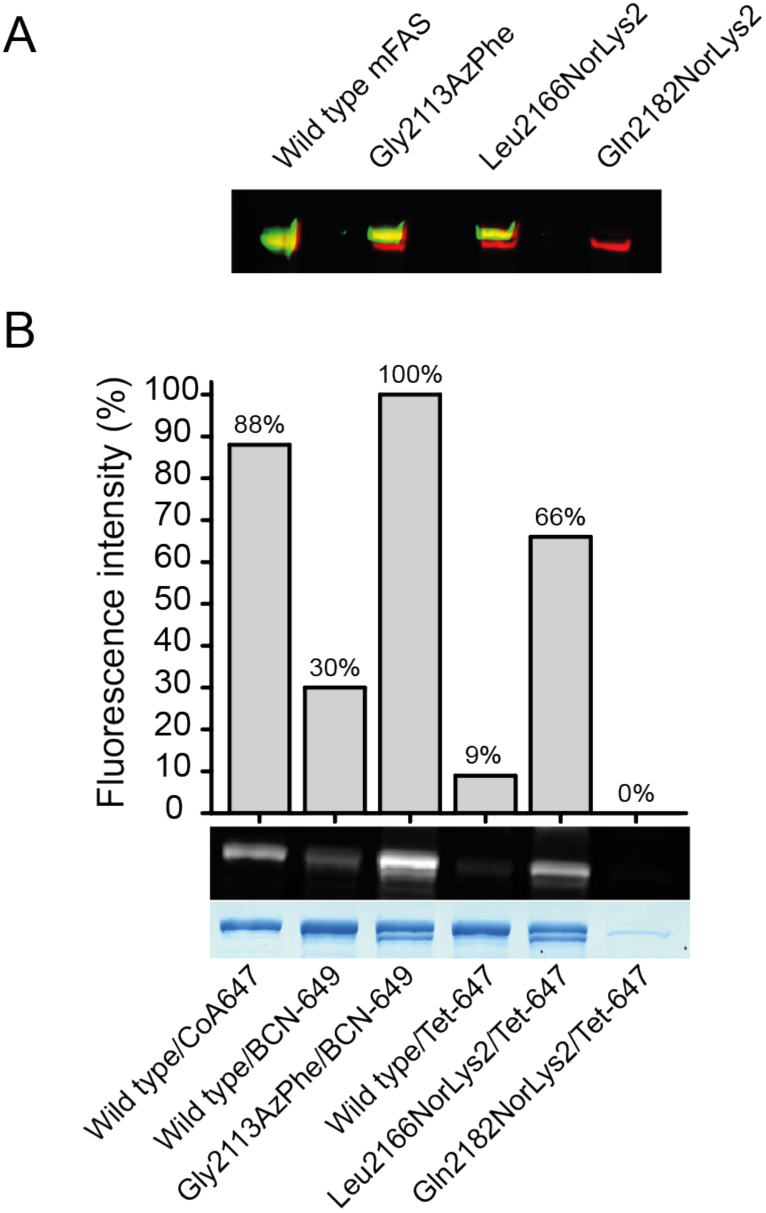
Production of ncAA-modified mFAS mutants for fluorescent labelling. A) Quantitative western blot of mFAS mutants. The red channel refers to antibodies conjugated with DyLight 755 against the N-terminal Strep-tag and the green channel to antibodies conjugated with DyLight 633 against the C-terminal His-tag, respectively. The missing green band in lane 4 indicates that expression of full-length Gln2182NorLys2 mFAS failed and that only a truncated construct without the C-terminal part was obtained. B) Fluorescent labelling of mFAS mutants. AzPhe mutants were labelled with BCN-POE_3_-NH-DY649P1 (BCN-649), NorLys2 mutants were labelled with 6-methyl-tetrazine-ATTO-647N (Tet-647) and the wild type mFAS was enzymatically modified at the ACP domain by a fluorescent CoA647-label by Sfp. DOL is determined by the relative in-gel fluorescence intensities at wavelength 650 nm related to the wild type reference. All fluorescence intensities were corrected by quantum efficiency of the respective fluorophore and correlated to the protein bands of the Coomassie-stained gel.

**Figure 7:**
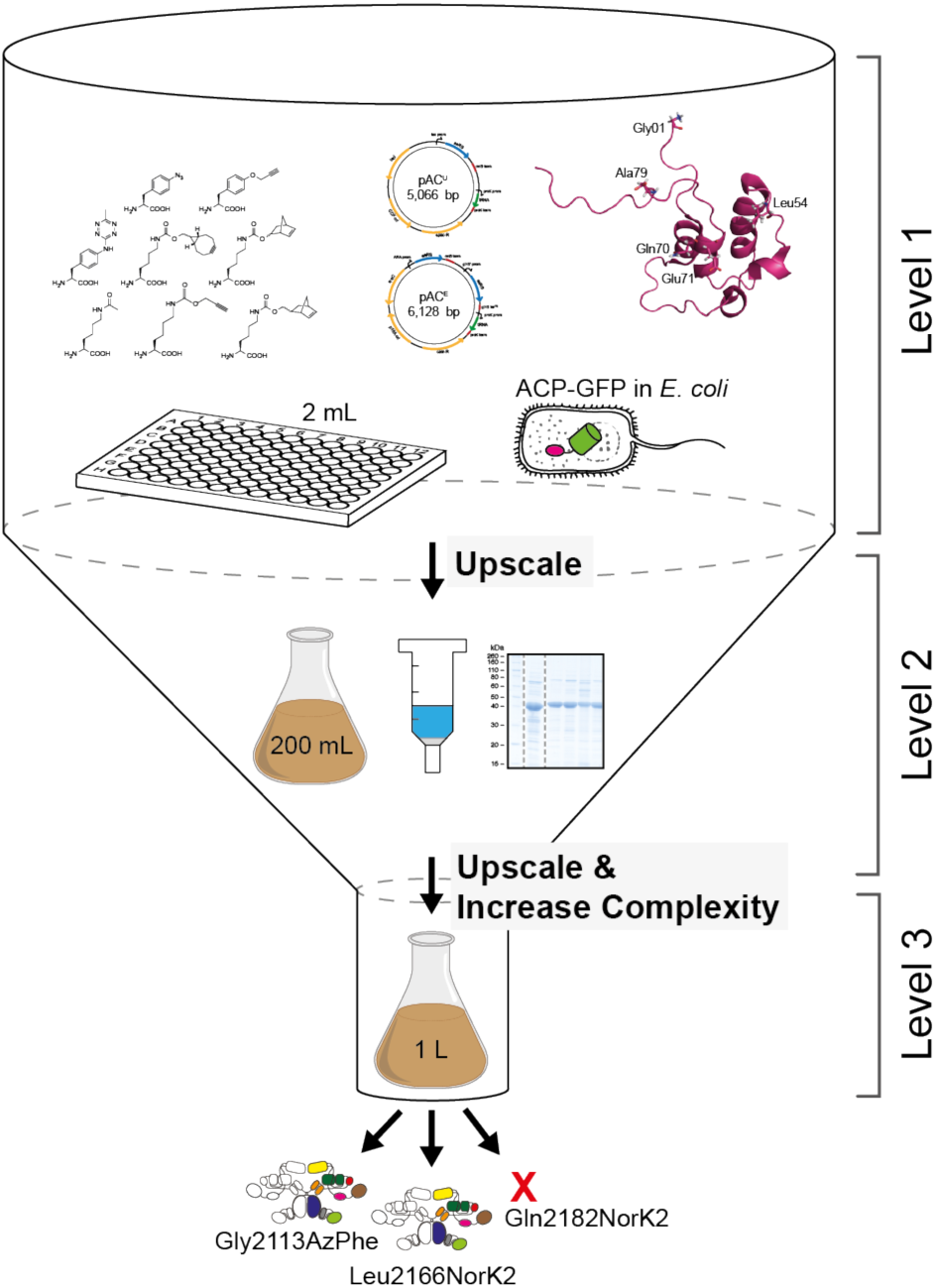
Overview of the workflow of this study. Workflow of amber codon suppression on mFAS divided into three different levels of project progress. Level 1 refers to the low-complex single-domain screening approach in 2 mL small scale cell cultures in 96-well plate format. GFP fluorescence is directly read out and serves as a measure for the efficiency of amber codon suppression. Level 2 refers to the upscaling of culture volumes to 200 mL using initial results from the reporter assay, which also allows obtaining purified protein for further analysis. In a final step, level 3 refers to the application of selected conditions and label positions, that were identified for an individual domain, for the full-length mFAS, being a representative for any comparable multidomain protein.

## Discussion

The last decades have shown amber codon suppression to be a powerful tool to investigate structure and function of proteins. The opportunity to label site-specifically with bioorthogonal functional groups has led to novel biophysical probes *in vitro* and *in vivo*^19,20,44–46^. So far, this technique has commonly been used to study small proteins, but has not been applied for large and complex multidomain systems.

Conformational variability is a fundamental property for the catalysis of many enzymes and especially megaenzymes as the animal FAS^6–8,13^. Here, we establish an efficient way to incorporate ncAAs site-specifically with subsequent labelling. Such probes make a range of biological applications accessible, such as photocrosslinking, electron paramagnetic resonance (EPR) spectroscopy, and FRET spectroscopy^44,47,48^. In case of animal FAS, labelled proteins would for example offer the prospect of addressing some of the fundamental questions to carrier domain-mediated substrate shuttling, i.e. towards the time scale of domain-domain interactions, or the influence of loaded intermediates on the mobility of the ACP domain.

Recently, we have established recombinant expression and purification of mFAS in *E. coli*^49^. Further, we have demonstrated that also parts and single domains of mFAS can be expressed individually, yielding for example the ACP domain as freestanding protein in high yields. Still incorporation of ncAAs by amber codon suppression into such a complex enzyme remains a challenging attempt, owing to the sensitivity of the protein to buffer and temperature changes as well as salt concentrations^50^. To circumvent high consumption of resources, we decided to approach this task by reducing complexity as far as possible. Specifically, we focused on the small ACP domain, and scaled down expression volumes to 2 mL cultures, which can be conveniently handled and analysed in 96-well format. In order to achieve a fast screening of amber codon suppression conditions, we fused the fluorescent protein GFP C-terminally to the ACP domain. This set-up allows to easily monitor the incorporation of the ncAA by the fluorescence emerging from the full-length fusion construct only.

Employing this set-up, we have compared two reported plasmid systems^27,28^, and tested the incorporation efficiency of eight different ncAAs, which predominantly allow copper-free click chemistry (Fig. 2A). For our set-up, the two suppressor plasmids pAC^U^ and pAC^E^, derived from pUltra and pEvol vectors, respectively, performed similarly well. The pAC^U^ vector has finally been used for its ease in protein production, as not requiring additional induction of the suppression system along with the protein of interest. In general, the TyrRS from *M. jannaschii* performed much better than the PylRS from *M. mazei* or *M. bakeri*, as the latter suffered from a high tendency to aggregate in *E. coli*^51,52^. As the two ncAAs AzPhe and NorLys2 showed optimal incorporation efficiencies of 26-35% and allow copper-free click chemistry, we chose to proceed further with these ncAAs incorporated with the pAC^U^ vector.

In addition to the choice of a suppression system, the reporter assay revealed that the site of ncAA incorporation in the ACP fold was critical for suppression efficiency^41^. Although all amber mutation sites were placed on the surface of the ACP domain, to avoid disturbing the protein fold or any protein-protein interactions in mFAS, both ncAAs were differently sensitive to incorporation sites. Whereas AzPhe was tolerated at all tested sites, NorLys2 was only introduced sufficiently into a disordered loop region. The higher tolerance to incorporate AzPhe at different positions may be explained by its smaller size preserving the integrity of the ACP fold.

Overall, the results of the upscale experiment and the reporter assay agree with one another. Incorporation efficiencies of the two different ncAAs at 5 different mutation sites were in line with data received from the reporter assay, with the only exception of NorLys2 incorporation at site Gly01. From SEC profiles of purified proteins, we were further able to conclude, that modification with ncAAs did not disturb the protein fold. With a drop in expression yield to 30–40% of the non-mutated reference construct, also the access to the protein remained satisfyingly high. As a quality control, we finally also employed the constructs in testing bioconjugation with fluorophores. Although there is a discrepancy between the DOL determined using relative in-gel fluorescence intensities and UV-Vis absorbance, both methods agreed that for our set-up the azido-BCN reaction is more efficient than the norbornene-tetrazine reaction. The labelling of the mFAS mutants Gly2113AzPhe and Leu2166NorLys2 finally led to two fluorescent probes, which can readily be subjected to fluorescence spectroscopic analysis^16^.

These results demonstrate the successful incorporation of ncAAs into a 540 kDa homodimer by amber codon suppression, with subsequent fluorescent labelling.

Such a systematic approach is necessary to tackle the challenges in application of amber codon suppression for multidomain proteins, comprising of a reporter assay on a single domain with upscaling of the culture volume and final modification of the full-length protein. Together, this procedure may be applied to any comparable biological system and can become a powerful tool to elucidate structure and conformational properties of multidomain proteins, as e.g. the homologous PKS family.

## Methods

### Cloning of suppressor plasmids pAC^U^ & pAC^E^

Suppressor plasmids pAC were constructed based on pUltra and pEVOL, as had been published by the lab of Schultz and coworkers^27,28^, by assembling cassettes amplified from the commercially available plasmids pCDF1b, pMAL-c5G and pEVOL_pBpF (was a gift from Peter Schultz (Addgene plasmid # 31190)). Phusion polymerase (Clontech) was used to generate PCR fragments, which were assembled with help of complementary primer overhang in a MegaPrimer PCR and subsequently cloned into the backbone using InFusion Cloning (Takara Bio). The pAC^U^ plasmids encode one copy of aaRS and suppressor tRNA under the tac promoter and rrnB terminator, and the proK promoter and proK terminator, respectively. The backbone of the pAC^U^ contains a CDF origin, spectinomycin resistance and *lacI* gene. The pAC^E^ plasmids encode two copies of aaRS, one under the arabinose promoter and the rrnB terminator, and one under the glnS’ promoter and glnS terminator, as well as one copy of suppressor tRNA under the proK promoter and proK terminator. The backbone of the pAC^E^ contains a p15A origin, chloramphenicol resistance and *araC* gene. Multiple point mutations were introduced to create a set of evolved aaRSs, specific for certain ncAAs. Three different sets of orthogonal suppressor pairs (aaRS/tRNA), derived from *M. jannaschii, M. mazei* and *M. barkeri*, are available. Genes of the orthogonal pair mmPylRS/tRNA were obtained from plasmid pJZ, which was a gift from Nediljko Budisa, and mbPylRS was obtained from pAcBac1.tR4-MbPyl, which was a gift from Peter Schultz (Addgene plasmid # 50832). Stellar cells were used for plasmid amplification. All mutations were confirmed by sequencing (Seqlab).

### Cloning of ACP-GFP-fusion constructs and amber mutations

The genes for ACP-GFP fusion constructs were cloned into a pET22b vector, which contains a pBR322 origin, ampicillin resistance and *lacI* gene. They are encoded under a T7 promoter and terminator, and feature a N-terminal Strep-tag and C-terminal His8-tag. The ACP-GFP construct termed wild type in this study contained a Met72Leu mutation, to prevent an alternative translation start and to reduce background GFP-fluorescence. An amber mutation was introduced site-specifically in the wild type gene and its position was varied throughout the ACP sequence to generate six different ACP-GFP mutant constructs, with different incorporation sites.

### General protein expression procedure

All constructs were transformed in *E. coli* BL21 Gold (DE3) cells (Agilent Technologies) following the provided protocol. For incorporation of ncAAs, plasmids encoding ACP-GFP constructs with amber mutations were co-transformed with the appropriate suppressor plasmid pAC^U^ or pAC^E^. LB agar (Lennox) transformation plates contained 1% glucose to suppress leaky expression, and were supplemented with either 100 µg/mL ampicillin for transformation of the ACP-GFP wild type, 50 µg/mL ampicillin and 25 µg/mL spectinomycin for co-transformation with pAC^U^ plasmids, or 50 µg/mL ampicillin and 17 µg/mL chloramphenicol for co-transformation with pAC^E^ plasmids. Colonies were grown at 37 °C overnight or at room temperature over weekend and stored at 4 °C up to several weeks. A randomly picked single clone was used to inoculate a pre-culture of Lysogeny Broth, supplemented with 1% glucose and respective antibiotics, which was grown at 37 °C and 180 rpm overnight. The pre-culture was used to inoculate Terrific Broth medium, supplemented with respective antibiotics. The cells were cultivated at 37 °C and 140–180 rpm until an OD_600_ of 0.5–0.7 was reached. The expression culture was supplemented with 2 mM final concentration of the ncAA and expression of the ACP-GFP constructs was induced with 0.25 mM final concentration of IPTG. Since the genes of pAC^U^ plasmids stand under a tac promoter, no additional induction was needed, whereas expression of the orthogonal suppressor pair from pAC^E^ plasmids was induced additionally with 0.02% final concentration of arabinose. Protein expression was carried out at 20 °C and 140–180 rpm overnight.

### Reporter assay

The reporter assay was performed in 2 mL scale in 96-well deep well plates in technical triplicates, using an ACP-GFP wild type construct without amber mutation as reference and negative samples of each construct without addition of ncAA. The cells were harvested by centrifugation (3,220 rcf for 5 min at 4 °C), washed and resuspended in 300 µL PBS.

### Expression of ACP-GFP constructs

Large scale expression of ACP-GFP constructs was carried out in 200 mL expression cultures. Prior to harvesting the cells by centrifugation (4,000 rcf for 20 min at 4 °C), 2 mL samples of cell cultures were taken for further quantification using GFP-fluorescence and western blot. All cell pellets were flash frozen in liquid nitrogen and stored at –80 °C until use.

### Expression of mFAS constructs

Large scale expression of mFAS constructs was carried out in 1 L expression cultures. The cells were harvested by centrifugation (4,000 rcf for 20 min at 4 °C) and subsequently purified.

### Purification of ACP-GFP constructs

The cell pellets were thawed on ice and resuspended in 10 mL His buffer (50 mM KPi, 200 mM NaCl, 20 mM imidazole, 10% glycerol, pH 7.4) containing DNase I and 1 mM EDTA. French pressure cell press was used for mechanical disruption at a pressure of 1000 bar and the cell debris was removed by centrifugation (50,000 rcf for 30 min at 4 °C). After addition of 2 mM MgCl_2_, the lysate was subjected to 3 mL (bead capacity 50 mg/mL) Ni-NTA Superflow resin (QIAGEN) and incubated for 1 h at 4 °C. Unbound protein was washed off with 5 column volumes His buffer and bound His-tagged protein was eluted with 2.5 column volumes His buffer containing 300 mM imidazole and additional 2 column volumes of His buffer with imidazole increased to 500 mM. The elution fractions were analysed by SDS-PAGE and size exclusion chromatography (SEC) over a Superdex 200 Increase 10/300 GL column (His buffer filtered and degassed). Protein samples were concentrated using an Amicon Ultra concentration device (Millipore), flash frozen in liquid nitrogen, and stored at -80 °C.

### Purification of ACP-GFP fluorophore conjugates

Excess fluorophore from bioconjugation reaction was removed by purification over 1 mg HisPur Ni-NTA Magnetic Beads (Thermo Fisher Scientific). At each purification step the beads were shortly vortexed, spun down and placed in a magnetic stand, so the liquid phase could be taken up with a pipette. The Ni-NTA beads were first equilibrated with 160 µL and additional 400 µL His buffer (50 mM KPi, 200 mM NaCl, 20 mM imidazole, 10% glycerol, pH 7.4). The bioconjugation reaction was diluted with one volume of His buffer and incubated with the Ni-NTA beads for 30 min in the dark on an end-over-end rotator. Unbound protein was washed off with two times 400 µL His buffer. In two elution steps, the bound His-tagged protein was incubated for 30 min, and 15 min respectively, in the dark on an end-over-end rotator with 50 µL His buffer containing 300 mM imidazole.

### Purification of mFAS constructs

The cell pellets were resuspended in 30 mL His buffer (50 mM NaPi, 450 mM NaCl, 10 mM imidazole, 20% glycerol, pH 7.6) containing DNase I and 1 mM EDTA. French pressure cell press was used for mechanical disruption at a pressure of 1000 bar and the cell debris was removed by centrifugation (50,000 rcf for 30 min at 4 °C). After addition of 2 mM MgCl_2_, the protein was bound to Ni-NTA resin (QIAGEN) and eluted at 300 mM imidazole. Additionally to Ni-IMAC the mFAS constructs were purified over a Strep-column (Iba), eluted with 2.5 mM desthiobiotin (Strep buffer: 250 mM KPi, 1 mM EDTA, 1 mM DTT, 10% glycerol, pH 7.4). Further purification was performed by size exclusion chromatography over a Superdex 200 Increase 10/300 GL column (Strep buffer filtered and degassed). Protein samples were concentrated using an Amicon Ultra concentration device (Millipore), flash frozen in liquid nitrogen, and stored at -80 °C.

### Quantification of GFP-fluorescence

The reporter assay samples and the 2 mL samples from large scale ACP-GFP expression were analysed by their GFP-fluorescence. An undiluted sample or 10– fold dilution of the resuspended cells in PBS was transferred into 96-well plates and OD_600_ and GFP fluorescence was measured at CLARIOstar (BMG). Blank corrected fluorescence values were normalized by OD_600_. Fluorescence intensity of the wild type was set to 100% and all other fluorescence signals were related to the wild type.

### Mass spectrometric protein analysis

Purified protein was analysed using nanoESI (Synapt G2-S) mass spectrometry. Protein buffer of the sample was changed to 0.1-1 M ammonium acetate in an Amicon Ultra concentration device (Millipore). Protein concentration of the samples was 1 mg/mL.

### Western blot analysis

From large scale mFAS expression cultures, 2 mL samples were taken for western blot analysis. OD_600_ from a 10–fold dilution was measured and the cells were normalized to an OD_600_ of 8. A small sample was analysed on a SDS-PAGE. The proteins were transferred from the analytical polyacrylamide gel onto a PVDF- membrane by an electrophoretic tank-blot method (25 V for 1 h). The membrane was subsequently blocked with 0.2% I-Block and 0.1% Tween-20 in PBS, treated first with monoclonal mouse anti-Strep antibody (StrepMAB classic, Iba) and monoclonal rabbit anti-His antibody (bethyl) (at 4 °C overnight), and secondly with IgG donkey anti-mouse DyLight 755 and IgG goat anti-rabbit DyLight 633 (Thermo Scientific) (light-protected at room temperature for 1 h). Excess antibodies were washed off in several washing steps with 0.2% I-Block and 0.1% Tween-20 in PBS. The western blot was developed at the Fusion SL Fluorescence Imaging System (Vilber Lourmat) with excitation in the near infrared to infrared range and using the emission filters F- 695 Y5 and F-820.

### Fluorescent bioconjugation

Bioconjugation of fluorophore (20–150 equiv.) and protein (1 equiv.) was performed in a copper-free environment at room temperature in the dark. The reaction proceeded in 25–100 µL His buffer for up to 4 hours. Protein mutants with the AzPhe were treated with the fluorophore BCN-POE_3_-NH-DY649P1, and mutants with the NorK2 were treated with 6-Methyl-tetrazine-ATTO-647N. Wild type ACP-GFP served as negative control, treated with the same fluorophores under the same conditions. As reference, 1 equiv. wild type ACP was phosphopantetheinylated with 5 equiv. of the fluorescent substrate CoA 647 (NEB), catalysed by 0.5 equiv. 4’- phosphopantetheinyl transferase Sfp from *B. subtilis*. The phosphopantetheinylation was performed in presence of 10 mM MgCl_2_ for 30–45 min at 37 °C in the dark. Subsequent to bioconjugation, an analytical SDS-PAGE was performed and fluorescent protein bands were detected at the Fusion SL Fluorescence Imaging System (Vilber Lourmat) with excitation in the near infrared range and using emission filter F-695 Y5. The FusionCapt Advance Solo 4 16.08a software was used to quantify the fluorescence of the protein bands on the polyacrylamide gel.

### UV-Vis spectra

UV-Vis spectra were recorded on a Carry 100 UV-Vis spectrophotometer (Agilent Technologies) from 800 nm to 220 nm wavelength in quartz glass cuvettes (50 µL sample). Excess fluorophore had been removed by purification over HisPur Ni-NTA Magnetic Beads (Thermo Fisher Scientific). The reference sample contained His buffer with 300 mM imidazole. Absorption at 650 nm (maximum absorption of the fluorophore) and at 485 nm (maximum absorption of GFP) was used to determine the degree of labelling, following equation 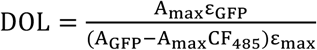, with ε_GFP_ and ε_max_ being the molar extinction coefficients of GFP and the fluorophores, respectively. The correction factor 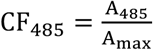 was determined from the absorption spectrum of the free fluorophore in water. All values for calculation are summarized in table 1.

**Table 1:** Calculation of the DOL with optical properties of used fluorophores

| Fluorophore | $A_{GFP}$ (mAU) | $\epsilon_{GFP}$ (L mol<sup>-1</sup> cm<sup>-1</sup>) | $A_{max}$ (mAU) | $\epsilon_{max}$ (L mol<sup>-1</sup> cm<sup>-1</sup>) | $CF_{485}$ |
| --- | --- | --- | --- | --- | --- |
| DY647P1 | 0.674271762 | 83300 | 1.49885743 | 250000 | 0.006 |
| DY649P1 | 0.454478949 | 83300 | 0.858465344 | 250000 | 0.004 |
| ATTO-647N | 0.479153782 | 83300 | 0.316676229 | 150000 | 0.013 |

## Data Availability

All data generated or analysed during this study are included in this published article and its Supplementary Information files. The plasmids (pAC^U^ and pAC^E^) generated during the current study are available from the corresponding author on reasonable request.

## Acknowledgements

We thank Khanh Vu Huu, Kudratullah Karimi and Prof. Nina Morgner for mass spectrometry analysis of proteins. We are also grateful to students Sina Manger and Vanessa Bause for assistance in the lab. We are thankful to Prof. Nediljko Budisa for supporting us at the beginning of the project. We would also like to thank Dr. Karthik Paithankar for proof-reading the manuscript.

## Author contribution

C.S.H. and A.R. performed molecular cloning of suppressor plasmids and plasmids encoding ACP-GFP and mFAS constructs. A.R. established expression and purification of mFAS. C.S.H. performed reporter assays, protein expression, purification experiments, fluorescent labelling and analysed corresponding data. B.G. and D.B. carried out experiments and assisted to establish the method under supervision of C.S.H. and A.R.. A.R. and B.G. synthesized ncAAs. A.R. conceived the project, which was further developed together with C.S.H.. M.G. designed the research and analysed data. C.S.H., A.R. and M.G. wrote the manuscript.

## Additional Information

### Supporting Information

accompanies this paper and is available online.

## References

1. Chang, S.-I. & Hammes, G. G. Structure and Mechanism of Action of a Multifunctional Enzyme: Fatty Acid Synthase. Acc. Chem. Res. 23, 363–369 (1990).

2. Maier, T., Leibundgut, M., Boehringer, D. & Ban, N. Structure and function of eukaryotic fatty acid synthases. Q. Rev. Biophys. 43, 373–422 (2010).

3. Barajas, J. F., Blake-Hedges, J. M., Bailey, C. B., Curran, S. & Keasling, J. D. Engineered polyketides: Synergy between protein and host level engineering. Synth. Syst. Biotechnol. 2, 147–166 (2017).

4. Hertweck, C. The Biosynthetic Logic of Polyketide Diversity. Angew. Chem. Int. Ed. 48, 4688–4716 (2009).

5. Smith, S. & Tsai, S.-C. The type I fatty acid and polyketide synthases: a tale of two megasynthases. Nat. Prod. Rep. 24, 1041–1072 (2007).

6. Maier, T., Leibundgut, M. & Ban, N. The Crystal Structure of a Mammalian Fatty Acid Synthase. Science 321, 1315–1322, doi:10.1126/science.1161269 (2008).

7. Viegas, M. F., Neves, R. P. P., Ramos, M. J. & Fernandes, P. A. Modeling of Human Fatty Acid Synthase and *in Silico* Docking of Acyl Carrier Protein Domain and Its Partner Catalytic Domains. J. Phys. Chem. B 122, 77–85 (2017).

8. Brignole, E. J., Smith, S. & Asturias, F. J. Conformational flexibility of metazoan fatty acid synthase enables catalysis. Nat. Struct. Mol. Biol. 16, 190–197, doi:10.1038/nsmb.1532 (2009).

9. Anselmi, C., Grininger, M., Gipson, P. & Faraldo-Gómez, J. D. Mechanism of Substrate Shuttling by the Acyl-Carrier Protein within the Fatty Acid Mega-Synthase. J. Am. Chem. Soc. 132, 12357–12364, doi:10.1021/ja103354w (2010).

10. Weissman, K. J. The structural biology of biosynthetic megaenzymes. Nat. Chem. Biol. 11, 660–670 (2015).

11. Rittner, A. & Grininger, M. Modular Polyketide Synthases (PKSs): A New Model Fits All? ChemBioChem 15, 2489–2493 (2014).

12. Jenni, S. et al. Structure of Fungal Fatty Acid Synthase and Implications for Iterative Substrate Shuttling. Science 316, 254–261 (2007).

13. Chang, S.-I. & Hammes, G. G. Amino Acid Sequences of Pyridoxal 5’-Phosphate Binding Sites and Fluorescence Resonance Energy Transfer in Chicken Liver Fatty Acid Synthase. Biochemistry 28, 3781–3788 (1989).

14. Yuan, Z. & Hammes, G. G. Fluorescence Studies of Chicken Liver Fatty Acid Synthase. J. Biol. Chem. 261, 13643–13651 (1986).

15. Benning, F. M. C. et al. High-Speed Atomic Force Microscopy Visualization of the Dynamics of the Multienzyme Fatty Acid Synthase. ACS Nano 11, 10852–10859, doi:10.1021/acsnano.7b04216 (2017).

16. Roy, R., Hohng, S. & Ha, T. A practical guide to single-molecule FRET. Nat Meth 5, 507–516 (2008).

17. Hohlbein, J., Gryte, K., Heilemann, M. & Kapanidis, A. N. Surfing on a new wave of single-molecule fluorescence methods. Phys. Biol. 7, 1–22 (2010).

18. Kim, Y. et al. Efficient Site-Specific Labeling of Proteins via Cysteines. Bioconjugate Chem. 19, 786–791 (2008).

19. Lang, K. & Chin, J. W. Cellular incorporation of unnatural amino acids and bioorthogonal labeling of proteins. Chem. Rev. 114, 4764–4806, doi:10.1021/cr400355w (2014).

20. Liu, C. C. & Schultz, P. G. Adding New Chemistries to the Genetic Code. Annu. Rev. Biochem. 79, 413–444, doi:10.1146/annurev.biochem.052308.105824 (2010).

21. Tsien, R. Y. The green fluorescent protein. Annu. Rev. Biochem. 67, 509–544 (1998).

22. Rostovtsev, V. V., Green, L. G., Fokin, V. V. & Sharpless, K. B. A stepwise huisgen cycloaddition process: copper(I)-catalyzed regioselective “ligation” of azides and terminal alkynes. Angew. Chem. Int. Ed. Engl. 41, 2596–2599 (2002).

23. Dommerholt, J. et al. Readily Accessible Bicyclononynes for Bioorthogonal Labeling and Three-Dimensional Imaging of Living Cells. Angew. Chem. Int. Ed. 49, 9422–9425, doi:10.1002/anie.201003761 (2010).

24. Lang, K. et al. Genetically encoded norbornene directs site-specific cellular protein labelling via a rapid bioorthogonal reaction. Nat. Chem. 4, 298–304, doi:10.1038/nchem.1250 (2012).

25. Hahn, A., Reschke, S., Leimkühler, S. & Risse, T. Ketoxime Coupling of *p*-Acetylphenylalanine at Neutral pH for Site-Directed Spin Labeling of Human Sulfite Oxidase. The Journal of Physical Chemistry B 118, 7077–7084, doi:10.1021/jp503471j (2014).

26. Choudhary, C. et al. Lysine Acetylation Targets Protein Complexes and Co-Regulates Major Cellular Functions. Science 325, 834–840 (2009).

27. Chatterjee, A., Sun, S. B., Furman, J. L., Xiao, H. & Schultz, P. G. A versatile platform for single- and multiple-unnatural amino acid mutagenesis in *Escherichia coli*. Biochemistry 52, 1828–1837, doi:10.1021/bi4000244 (2013).

28. Young, T. S., Ahmad, I., Yin, J. A. & Schultz, P. G. An Enhanced System for Unnatural Amino Acid Mutagenesis in *E. coli*. J. Mol. Biol. 395, 361–374, doi:10.1016/j.jmb.2009.10.030 (2010).

29. Chin, J. W. et al. Addition of *p*-Azido-l-phenylalanine to the Genetic Code of *Escherichia coli*. J. Am. Chem. Soc. 124, 9026–9027, doi:10.1021/ja027007w (2002).

30. Kaya, E. et al. A Genetically Encoded Norbornene Amino Acid for the Mild and Selective Modification of Proteins in a Copper-Free Click Reaction. Angew. Chem. Int. Ed. 51, 4466–4469, doi:10.1002/anie.201109252 (2012).

31. Schneider, S. et al. Structural Insights into Incorporation of Norbornene Amino Acids for Click Modification of Proteins. ChemBioChem 14, 2114–2118, doi:10.1002/cbic.201300435 (2013).

32. Borrmann, A. et al. Genetic Encoding of a Bicyclo[6.1.0]nonyne-Charged Amino Acid Enables Fast Cellular Protein Imaging by Metal-Free Ligation. ChemBioChem 13, 2094–2099 (2012).

33. Deiters, A. & Schultz, P. G. *In vivo* incorporation of an alkyne into proteins in *Escherichia coli*. Bioorg. Med. Chem. Lett. 15, 1521–1524, doi:10.1016/j.bmcl.2004.12.065 (2005).

34. Seitchik, J. L. et al. Genetically Encoded Tetrazine Amino Acid Directs Rapid Site-Specific *in Vivo* Bioorthogonal Ligation with *trans*-Cyclooctenes. J. Am. Chem. Soc. 134, 2898–2901, doi:10.1021/ja2109745 (2012).

35. Nguyen, D. P. et al. Genetic Encoding and Labeling of Aliphatic Azides and Alkynes in Recombinant Proteins *via* a Pyrrolysyl-tRNA Synthetase/tRNACUA Pair and Click Chemistry. J. Am. Chem. Soc. 131, 8720–8721 (2009).

36. Umehara, T. et al. N-Acetyl lysyl-tRNA synthetases evolved by a CcdB-based selection possess N-acetyl lysine specificity *in vitro* and *in vivo*. FEBS Lett. 586, 729–733, doi:10.1016/j.febslet.2012.01.029 (2012).

37. Kobayashi, T. et al. Structural basis for orthogonal tRNA specificities of tyrosyl-tRNA synthetases for genetic code expansion. Nat. Struct. Biol. 10, 425–432 (2003).

38. Wang, Y.-S., Fang, X., Wallace, A. L., Wu, B. & Liu, W. R. A Rationally Designed Pyrrolysyl-tRNA Synthetase Mutant with a Broad Substrate Spectrum. J. Am. Chem. Soc. 134, 2950–2953, doi:10.1021/ja211972x (2012).

39. Young, D. D. et al. An Evolved Aminoacyl-tRNA Synthetase with Atypical PolysubstrateSpecificity. Biochemistry 50, 1894–1900, doi:10.1021/bi101929e (2011).

40. Schultz, K. C. et al. A Genetically Encoded Infrared Probe. J. Am. Chem. Soc. 128, 13984–13985, doi:10.1021/ja0636690 (2006).

41. Pott, M., Schmidt, M. J. & Summerer, D. Evolved Sequence Contexts for Highly Efficient Amber Suppression with Noncanonical Amino Acids. ACS Chem. Biol. 9, 2815–2822 (2014).

42. Quadri, L. E. N. et al. Characterization of Sfp, a *Bacillus subtilis* Phosphopantetheinyl Transferase for Peptidyl Carrier Protein Domains in Peptide Synthetases. Biochemistry 37, 1585–1595 (1998).

43. Lee, K. K., Da Silva, N. A. & Kealey, J. T. Determination of the extent of phosphopantetheinylation of polyketide synthases expressed in *Escherichia coli* and *Saccharomyces cerevisiae*. Anal. Biochem. 394, 75–80, doi:10.1016/j.ab.2009.07.010 (2009).

44. Tyagi, S. & Lemke, E. A. Single-molecule FRET and crosslinking studies in structural biology enabled by noncanonical amino acids. Curr. Opin. Struct. Biol. 32, 66–73, doi:10.1016/j.sbi.2015.02.009 (2015).

45. Chin, J. W. Expanding and Reprogramming the Genetic Code of Cells and Animals. Annu. Rev. Biochem. 83, 379–408, doi:10.1146/annurev-biochem-060713-035737 (2014).

46. Kim, C. H., Axup, J. Y. & Schultz, P. G. Protein conjugation with genetically encoded unnatural amino acids. Curr. Opin. Chem. Biol. 17, 412–419, doi:10.1016/j.cbpa.2013.04.017 (2013).

47. Hubbell, W. L., Cafiso, D. S. & Altenbach, C. Identifying conformational changes with site-directed spin labeling. 7, 735–739 (2000).

48. Schmidt, M. J. et al. EPR Distance Measurements in Native Proteins with Genetically Encoded Spin Labels. ACS Chem. Biol. 10, 2764–2771, doi:10.1021/acschembio.5b00512 (2015).

49. Rittner, A., Paithankar, K. S., Huu, K. V. & Grininger, M. Characterization of the Polyspecific Transferase of Murine Type I Fatty Acid Synthase (FAS) and Implications for Polyketide Synthase (PKS) Engineering. ACS Chem. Biol., doi:10.1021/acschembio.7b00718 (2018).

50. Rangan, V. S., Joshi, A. K. & Smith, S. Mapping the Functional Topology of the Animal Fatty Acid Synthase by Mutant Complementation *in Vitro*. Biochemistry 40, 10792–10799, doi:10.1021/bi015535z (2001).

51. Wan, W., Tharp, J. M. & Liu, W. R. Pyrrolysyl-tRNA synthetase: An ordinary enzyme but an outstanding genetic code expansion tool. Biochim. Biophys. Acta 1844, 1059–1070, doi:10.1016/j.bbapap.2014.03.002 (2014).

52. Jiang, R. & Krzycki, J. A. PylSn and the homologous N-terminal domain of pyrrolysyl-tRNA synthetase bind the tRNA that is essential for the genetic encoding of pyrrolysine. J. Biol. Chem. 287, 32738–32746, doi:10.1074/jbc.M112.396754 (2012).

