## Supplementary Materials for "Site-Specific Labelling of Multidomain Proteins by Amber Codon Suppression"

##### Table of contents:

**Supplementary Methods:**

1. Synthesis of non-canonical amino acids
2. Detailed cloning procedure

**Supplementary Data:**

Mass spectrometric analysis of ACP-GFP constructs

**Supplementary Figures:**

Figure S1: Cloning scheme of $pAC^U$ and $pAC^E$ vectors

Figure S2: Screening for the most efficient suppression vector system

Figure S3: Testing the effect of different ncAA concentrations

Figure S4: Optimizing the click reaction conditions for fluorescent labelling

Figure S5: Generation of ncAA-modified mFAS mutants for fluorescent labelling

Figure S6: Large scale expression and purification of ACP-GFP mutants

Figure S7: Fluorescent labelling of ACP-GFP mutants

**Supplementary Tables**

Table S1: List of primers

Table S2: List of amber codon suppressor plasmids

### Supplementary Methods

#### 1. Synthesis of non-canonical amino acids

All chemicals were used as purchased from Sigma-Aldrich without further processing. Protected amino acids were ordered at Iris Biotech. All syntheses except for AzPhe were performed under dry conditions and argon atmosphere.

##### 1.1. Synthesis of AzPhe (**2**)[1]

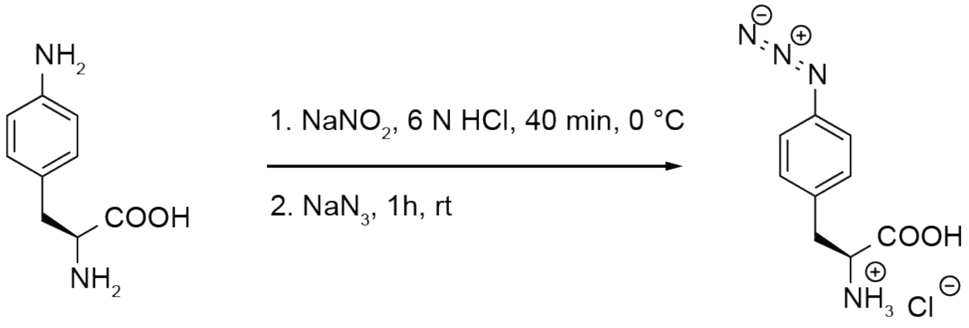


4-Aminophenylalanine **1** (10.0 g, 55.4 mmol, 1.0 equiv.) was dissolved in 6 N HCl solution (200 mL) and cooled down to 0 °C. Subsequently, a solution of $NaNO_2$ (5.0 g, 72 mmol, 1.3 equiv.) in water (20 mL) was prepared and slowly added. The reaction mixture was stirred for 40 min at 0 °C. Then, a solution of $NaN_3$ (5.4 g, 83.2 mmol, 1.5 equiv.) in water (20 mL) was prepared and added in one portion. The reaction mixture was kept at 0 °C for another 5 min and allowed to warm to room temperature for 1 h. After evaporation of the solvent *in vacuo*, the crude product was purified on silica gel (7:3 DCM/MeOH, $R_f$ = 0.25), affording **2** as slightly yellow solid (11.5 g, 85%).

$^1$H-NMR (250 MHz, DMSO-$d_6$) δ (ppm) = 7.34 (d, $^3J$ = 8.4 Hz, 2H), 7.10 (d, $^3J$ = 8.4 Hz, 2H), 4.13 (dd, $^3J$ = 6.4 Hz, 6.3 Hz, 1H), 3.17-3.03 (m, 2H).

IR (KBr): Azide band at 2140 $cm^{-1}$ matches with the literature[2].

#### 1.2. Synthesis of PrPhe (**6**)

##### 1.2.1. Synthesis of methyl-2-(*tert*-butoxycarbonyl)amino)-3-(4-(ethynyloxy)phenyl)propanoate **4**[3]

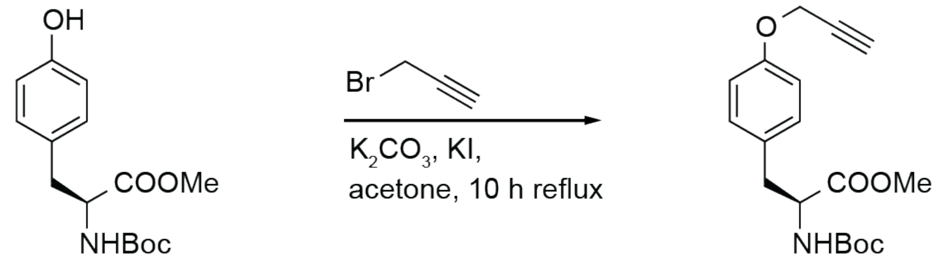


**3** **4**

$K_2CO_3$ (9.10 g, 67.5 mmol, 1.5 equiv.) and KI (0.72 g, 4.50 mmol, 0.1 equiv.) were added to a stirred solution of Boc-L-Tyr-OMe **3** (13.1 g, 45.0 mmol, 1.0 equiv.) in acetone (150 mL) at room temperature. Propargyl bromide (6 mL, 45 mmol, 1 equiv.) was added dropwise over 30 min and the resulting mixture was heated under reflux for 10 h. The reaction mixture was cooled on ice and filtered two times. The filtrate was concentrated *in vacuo* and the residue was re-dissolved in ethyl acetate (100 mL), washed twice with water (30 mL), and dried over $MgSO_4$. Evaporation of the solvent *in vacuo* afforded **4** as yellow oil (14.1 g, 94%).

$^1$H-NMR (300 MHz, $CDCl_3$): δ (ppm) = 7.06 (d, $^3J$ = 8.75 Hz, 2H), 6.91 (d, $^3J$ = 8.75 Hz, 2H), 4.97-4.96 (m, 1H), 4.67 (d, $^3J$ = 2.50 Hz, 2H), 4.54-4.53 (m, 1H), 3.71 (s, 3H), 3.11-3.03 (m, 1H), 2.53-2.52 (m, 1H), 1.42 (s, 9H).

1.2.2. Synthesis of 2-((*tert*-butoxycarbonyl)amino)-3-(4-(ethynyloxy)phenyl)propanoic acid **5**[4]

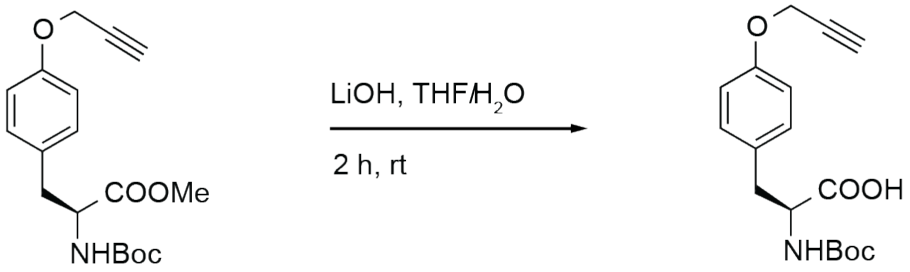


**4** **5**

The propargyl derivative Boc-L-PrPhe-OMe **4** (13.3 g, 40.0 mmol, 1.0 equiv.) was dissolved in THF (100 mL) and 1 M LiOH solution (85 mL, 85 mmol, 2.0 equiv.) was added carefully to the stirred solution at room temperature. After 1 h, completion of the reaction was checked by TLC (1:1 EtOAC/n-Hexan, $R_f = 0.8$). The reaction mixture was acidified with conc. HCl (10 mL) and the two phases were separated. The aqueous phase was extracted three times with ethyl acetate and the combined organic phases were washed with brine, and dried over $MgSO_4$. Evaporation of the solvent *in vacuo* afforded **5** as yellow oil (11.7 g, 91%).

$^1$H-NMR (250 MHz, $CDCl_3$): δ (ppm) = 7.13 (d, $^3J$ = 8.51 Hz, 2H), 6.92 (d, $^3J$ = 8.70 Hz, 2H), 4.68 (d, $^3J$ = 2.40 Hz, 2H), 3.19-2.90 (m, 2H), 2.52 (t, $^3J$ = 2.40 Hz, 1H), 2.05 (s, 1H), 1.43 (s, 9H).

1.2.3. Synthesis of 2-amino-3-(4-(ethynyloxy)phenyl)propanoic acid hydrochloride **6**[5]

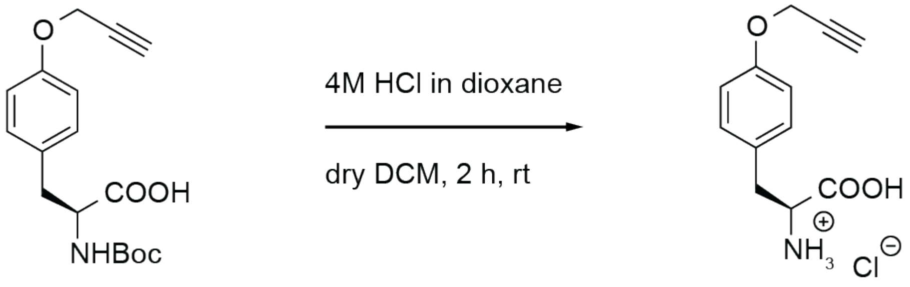


**5** **6**

The carboxylic acid Boc-L-PrPhe **5** (11.7 g, 37.0 mmol, 1.0 equiv.) was dissolved in DCM (25 mL) and 4 M HCl in dioxane (15 mL, 60 mmol, 1.6 equiv.) were added to the stirred solution at room temperature. After 2.5 h of reaction, the solvent was evaporated *in vacuo* to afford **6** as colourless solid (9.33 g, 99%).

$^1$H-NMR (400 MHz, DMSO-d$_6$): δ (ppm) = 8.40 (br s, 2H), 7.21 (d, $^3J$ = 8.56 Hz, 2H), 6.94 (d, $^3J$ = 8.56 Hz, 2H), 4.77 (d, $^3J$ = 2.32 Hz, 2H), 4.11-4.08 (m, 1H), 3.57-3.56 (m, 1H), 3.08 (d, $^3J$ = 6.11 Hz, 2H).

MALDI-MS (m/z): [M+Na]$^+$ calcd for $C_{12}H_{13}NO_3$ 242.07876, meas. 242.07914.

#### 1.3. Synthesis of TetPhe (**12**)

##### 1.3.1. Synthesis of methylthiocarbohydrazide **8**[5]

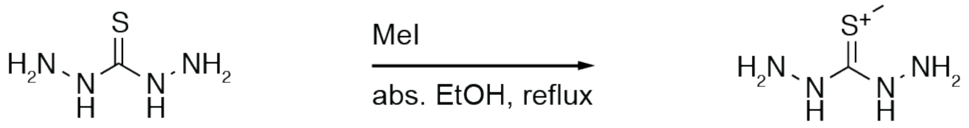


Thiocarbohydrazide **7** (24.1 g, 0.23 mol, 1.0 equiv.) was suspended in abs. ethanol (800 mL) and heated under reflux. Subsequently, iodomethane (16 mL, 0.3 mol, 1.1 equiv.) was added and heating was continued for 1 h. The solution was filtered and stirred for 1 h at room temperature, then kept at 4 °C overnight. The solvent was removed by decanting and remaining solvent was evaporated *in vacuo* to afford **8** as colourless needles (33.3 g, 59%).

$^{1}$H-NMR (250 MHz, DMSO-d$_6$): δ (ppm) = 2.38 (s, 3H).

##### 1.3.2. Synthesis of 3-methyl-6-(methylthio)-1,2,4,5-tetrazine **9**[6]

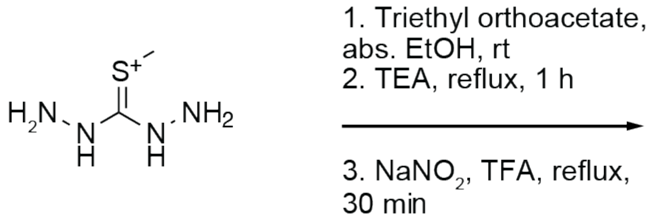


Triethyl orthoacetate (11 mL, 62 mmol, 1.4 equiv.) was added to a stirred suspension of methylthiocarbohydrazide **8** (11.2 g, 45.1 mmol, 1.0 equiv.) in abs. ethanol (300 mL) at room temperature. After 30 min, triethylamine (6.3 mL, 45 mmol, 1.0 equiv.) was added and the mixture was heated under reflux for 1 h. Subsequently, $NaNO_2$ (6.40 g, 92.8 mmol, 2.1 equiv.) and TFA (3.5 mL, 45 mmol, 1.0 equiv.) were added and heating was continued for 30 min. After addition of *n*-hexane (300 mL), the solution was degassed with Ar and stirred for 30 min at room temperature. Then, water (600 mL) was added, and the solution was extracted three times with diethyl ether (300 mL), and dried over $MgSO_4$. The solvent was evaporated *in vacuo*, and the crude product was purified on silica gel (19:1 *n*-hexane/DCM) to afford **9** as red oil (1.50 g, 23%).

$^{1}$H-NMR (250 MHz, $CDCl_3$): δ (ppm) 2.97 (s, 3H), 2.72 (s, 3H).

1.3.3. Synthesis of 2-((*tert*-butoxycarbonyl)amino)-3-(4-((6-methyl-1,2,4,5,-tetrazine-3-yl)amino)phenyl)propanoic acid **11**[5]

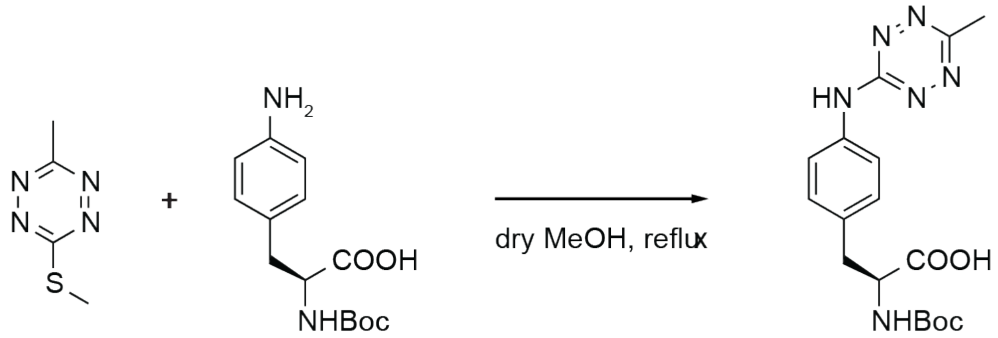


**9** **10** **11**

The tetrazine derivative **9** (0.57 g, 4.00 mmol, 1.1 equiv.) and Boc-D-4-aminophenylalanine **10** (1.02 g, 3.60 mmol, 1.0 equiv.) were suspended in methanol (10 mL) and heated under reflux for 18 h. The solvent was evaporated *in vacuo,* and the crude product was purified on silica gel (95:5 $CHCl_3$/methanol) to afford **11** as orange crystals (0.36 g, 23%).

$^{1}$H-NMR (250 MHz, DMSO-$d_6$): δ (ppm) = 12.60 (s, 1H), 10.62 (s, 1H), 7.63 (d, $^{3}J$ = 7.5 Hz, 2H), 7.25 (d, $^{3}J$ = 7.5 Hz, 2H), 4.08-4.05 (m, 1H), 3.04-2.79 (m, 2H), 2.78 (s, 3H), 1.34 (s, 9H).

1.3.4. Synthesis of 2-amino-3-(4-((6-methyl-1,2,4,5-tetrazine-3-yl)amino)phenyl)propanoic acid hydrochloride **12**[5]

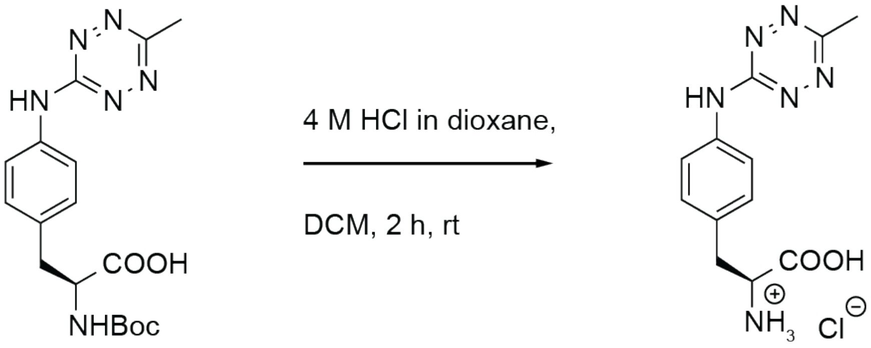


**11** **12**

The Boc-protected tetrazine derivative **11** (225 mg, 1.58 mmol, 1.0 equiv.) was dissolved in DCM (8 mL), and 4 M HCl in dioxane (8.0 mL, 32 mmol, 20 equiv.) was added. The solution was stirred for 2 h at room temperature. The solvent was evaporated *in vacuo* to afford **12** as red crystals (161 mg, 87%).

$^1$H-NMR (400 MHz, DMSO-d$_6$): δ (ppm) = 10.70 (s, 1H), 8.41 (br s, 2H), 7.67 (d, $^3J$ = 8.56 Hz, 2H), 7.28 (d, $^3J$ = 8.56 Hz, 2H), 4.14 (br t, $^3J$ = 6.30 Hz, 1H), 3.11 (d, $^3J$ = 6.30 Hz, 2H), 2.78 (s, 3H).

MALDI-MS (m/z): $[M+Na]^+$ calcd for $C_{12}H_{14}N_6O_2$ 297.10704, meas. 297.10681.

#### 1.4. Synthesis of AcLys (**15**)

##### 1.4.1. Synthesis of 6-acetamido2-((*tert*-butoxycarbonyl)amino)hexanoic acid **14**

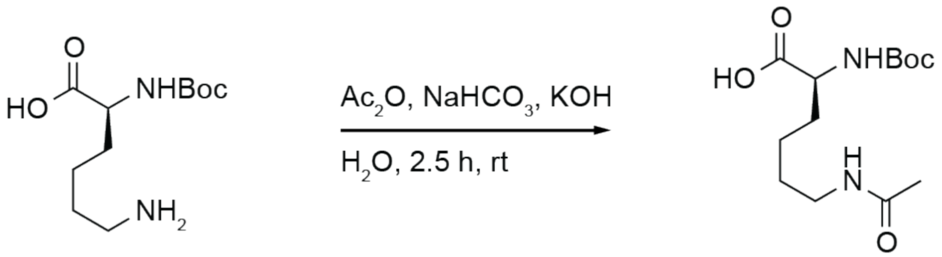


**13** **14**

To a solution of Boc-L-lysine **13** (3.06 g, 12.2 mmol, 1.0 equiv.) in water (60 mL), $NaHCO_3$ (3.12 g, 36.6 mmol, 3.0 equiv.) and KOH (0.85 g, 12.2 mmol, 1.0 equiv.) were added. Subsequently, acetic anhydride (1.2 mL, 12 mmol, 1.0 equiv.) was added dropwise over 10 min and the solution was stirred for 2.5 h at room temperature. After addition of ethyl acetate (60 mL), the aqueous phase was acidified with conc. HCl to pH 2. The two phases were separated, and the aqueous phase was extracted three times with ethyl acetate (30 mL). The organic phases were combined, washed with brine and dried over $MgSO_4$. Evaporation of the solvent *in vacuo* afforded **14** as slightly yellow oil, which was used directly in the next step without further purification.

$^1$H-NMR (250 MHz, DMSO-d$_6$) δ (ppm) = 12.11 (s, 2H), 7.79 (t, $^3J$ = 5.4 Hz, 1H), 7.01 (d, $^3J$ = 8.2 Hz, 1H), 3.89-3.77 (m, 1H), 3.00 (q, $^3J$ = 5.8 Hz, 2H), 2.10 (s, 2H), 1.78 (s, 2H), 1.71-1.47 (m, 2H), 1.39 (s, 9H).

1.4.2. Synthesis of 6-acetamido-2-aminohexanoic acid hydrochloride **15**[5]

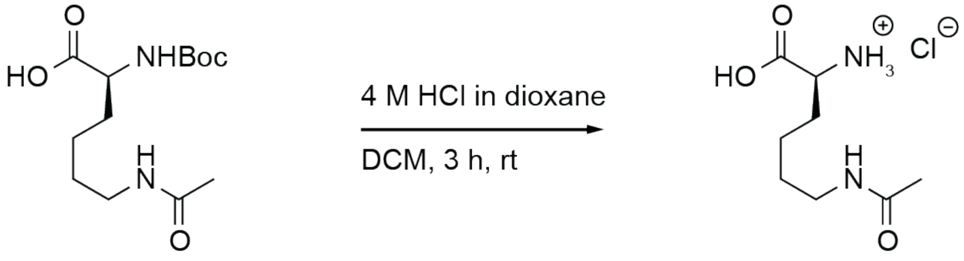


**14** **15**

A solution of 6-acetylamino-Boc-L-lysine **14** in dioxane (38 mL) and 4 M HCl in dioxane (3 mL, 12 mmol, 1.0 equiv.) was stirred for 3 h at room temperature. After evaporation of the solvent *in vacuo*, the residue was re-dissolved in 1 M HCl (60 mL). Subsequently, the solvent was removed azeotrope *in vacuo* by addition of toluene (4×25 mL) to afford a yellow oil. Lyophilisation obtained **15** as an off-white powder (2.75 g, 100%).

$^1$H-NMR (250 MHz, DMSO-d$_6$) δ (ppm) = 8.54-8.47 (m, 2H), 8.06-8.03 (m, 1H), 3.81 (q, $^3J$ = 5.5 Hz, 1H), 3.03-2.96 (m, 2H), 1.79 (s, 5H), 1.45-1.27 (m, 4H).

ESI-MS (m/z): $[M+H]^+$ calcd for $C_8H_{16}N_2O_3$ 189.12, meas. 189.12.

#### 1.5. Synthesis of PrLys (**17)**

##### 1.5.1. Synthesis of 2-((*tert*-butoxycarbonyl)amino)-6-(((prop-2-yn-1-yloxy)carbonyl)amino)hexanoic acid **16**[7]

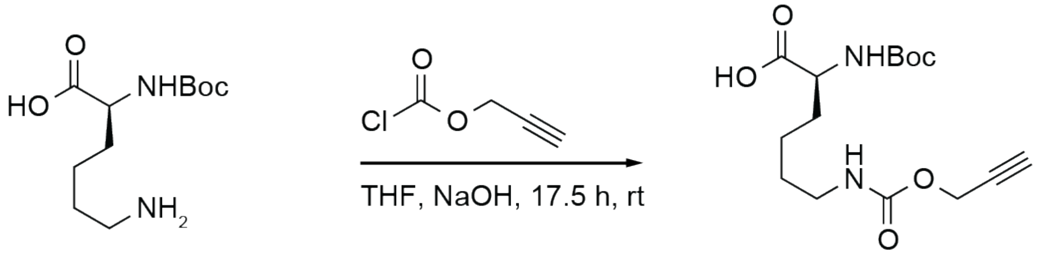


**13** **16**

A solution of Boc-L-lysine **13** (3.00 g, 12.2 mmol, 1.3 equiv.) in THF (30 mL) and 1 M NaOH (30 mL) was cooled down to 0 °C. Subsequently, propargyl chloroformate (1.0 mL, 9.7 mmol, 1.0 equiv.) was added dropwise over 10 min. The solution was stirred at room temperature for 17.5 h. After addition of ethyl acetate (60 mL), the aqueous phase was acidified with conc. HCl to pH 1.5. The two phases were separated, and the aqueous phase was extracted ethyl acetate (30 mL). The organic phases were combined, washed with brine and dried over $MgSO_4$. Evaporation of the solvent *in vacuo* afforded **16** as yellow oil, which was used directly in the next step without further purification.

$^1$H-NMR (400 MHz, DMSO-d$_6$): δ (ppm) = 12.38 (s, 1H), 7.30 (t, $^3J$ = 4.3 Hz, 1H), 7.00 (d, $^3J$ = 7.7 Hz, 1H), 4.80 (d, $^3J$ = 2.4 Hz, 1H), 4.59 (d, $^3J$ = 2.2 Hz, 2H), 3.86-3.77 (m, 1H), 3.46-3.43 (m, 1H), 2.96 (q, $^3J$ = 5.9 Hz, 2H), 1.38 (s, 9H), 1.68-1.46 (m, 2H), 1.33-1.24 (m, 2H).

1.5.2. Synthesis of 2-amino-6-(((prop-2-yn-1-yloxy)carbonyl)amino)hexanoic acid hydrochloride **17**[5]

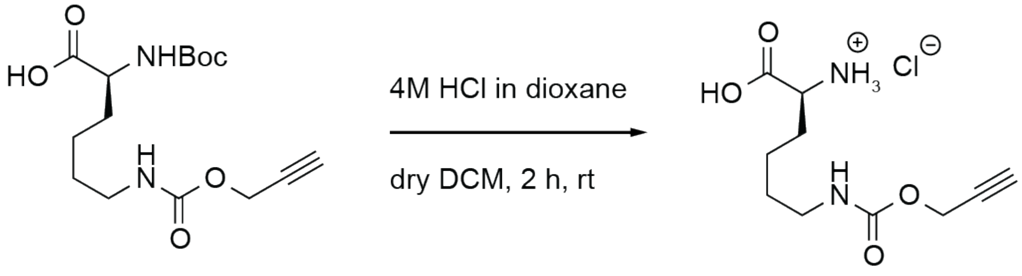


**16** **17**

A solution of 6-propargylformylamino-Boc-L-lysine **16** in dioxane (30 mL) and 4 M HCl in dioxane (2.4 mL, 9.6 mmol, 0.8 equiv.) was stirred for 4 h at room temperature. After evaporation of the solvent *in vacuo*, the residue was re-dissolved in 1 M HCl (80 mL). Subsequently, the solvent was removed azeotrope *in vacuo* by three times addition of toluene (25 mL) to afford **17** as an off-white powder (1.70 g, 55.0%).

$^1$H-NMR (400 MHz, DMSO-$d_6$): δ (ppm) = 8.40 (br s, 2H), 7.36 (t, $^3J$ = 5.44 Hz, 1H), 4.60 (d, $^3J$ = 2.32 Hz, 2H), 3.85-3.84 (m, 1H), 3.48 (t, $^3J$ = 2.32 Hz, 1H), 2.98 (q, $^3J$ = 6.05 Hz, 2H), 1.79-1.78 (m, 2H), 1.41-1.38 (m, 3H).

MALDI-MS (m/z): $[M+Na]^+$ calcd for $C_{10}H_{16}N_2O_4$ 251.10023, meas. 251.10030.

#### 1.6. Synthesis of NorLys1 (**22)**

##### 1.6.1. Synthesis of bicyclo[2.2.1]hept-5-en-2-ol **19**[8]

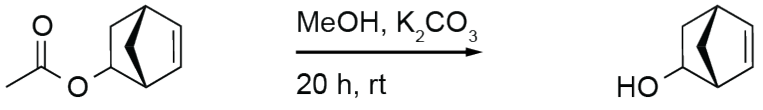


**18** **19**

A mixture of 5-norbornene-2-yl acetate **18** (10.0 g, 65.7 mmol, 1.0 equiv.) and $K_2CO_3$ (9.60 g, 69.7 mmol, 1.1 equiv.) in methanol (100 mL) was stirred for 20 h at room temperature. After evaporation of the solvent *in vacuo*, the residue was re-dissolved in water (300 mL), and extracted with ethyl acetate (5×100 mL). Evaporation of the solvent *in vacuo* afforded **19** as off-white crystals (6.20 g, 85.7%).

$^1$H-NMR (400 MHz, $CDCl_3$) δ (ppm) = 6.46-6.45 (m, 1H), 6.07-6.06 (m, 1H), 4.48-4.47 (m, 1H), 3.00 (s, 1H), 2.82 (s, 1H), 2.14-2.10 (m, 1H), 1.33-1.28 (m, 3H).

##### 1.6.2. Synthesis of bicyclo[2.2.1]hept-5-en-2-yl 2-(2,5-dioxopyrrolidin-1-yl)acetate **20**[9]

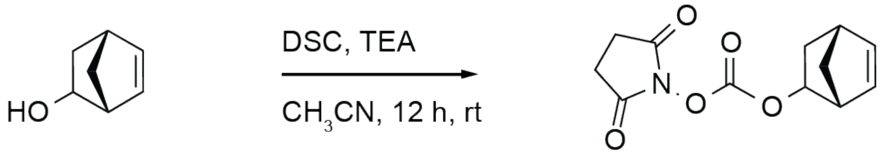


**19** **20**

To a solution of the norbornene derivative **19** (3.0 g, 27 mmol, 1.0 equiv.) in acetonitrile (100 mL), triethylamine (12 mL, 87 mmol, 3.2 equiv.) and *N*,*N*'-disuccinimidyl carbonate (11.3 g, 44.0 mmol, 1.2 equiv.) were added, and the mixture was stirred for 12 h at room temperature. After evaporation of the solvent *in vacuo*, the crude product was purified on silica gel (99.5:0.5 DCM/$Et_2O$) to afford **20** as slightly yellow crystals (6.20 g, 92.0%).

$^1$H-NMR (250 MHz, $CDCl_3$) δ (ppm) = 6.41-6.30 (m, 1H), 6.04-5.94 (m, 1H), 5.36 and 4.73 (m, 1H), 3.27 and 3.08 (m, 1H), 2.91 (s, 1H), 2.82 (s, 4H), 2.24-2.14 (m, 1H), 1.68-1.10 (m, 3H).

1.6.3. Synthesis of 6-(((bicyclo[2.2.1]hept-5-en-2-yloxy)carbonyl)amino)-2-((tert-butoxycarbonyl)amino)hexanoic acid **21**[9]

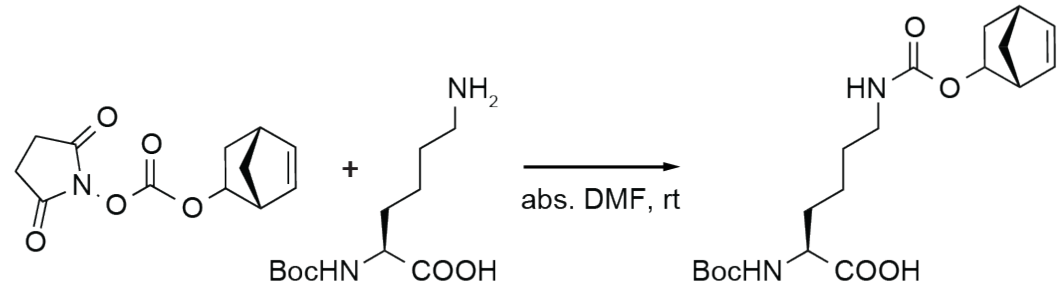


**20** **13** **21**

To a solution of the activated norbornene **20** (2.50 g, 10.0 mmol, 1.0 equiv.) in DMF (35 mL), the Boc-protected amino acid Boc-Lys-OH **13** (3.20 g, 13.0 mmol, 1.3 equiv.) was added, and the mixture was stirred over night at room temperature. After addition of water (300 mL), the mixture was extracted with ethyl acetate (150 mL). The organic phases were combined, washed with water (150 mL) and once with brine (75 mL), and dried over $MgSO_4$. Evaporation of the solvent *in vacuo* afforded **21** as white foam, which was used directly in the next step without further purification.

$^1$H-NMR (400 MHz, $CDCl_3$) δ (ppm) = 6.29-6.19 (m, 1H), 5.95-5.93 (m, 1H), 5.30-5.22 (m, 2H), 4.93-4.09 (m, 2H), 3.11 (br s, 2H), 2.80-2.79 (m, 1H), 2.09-2.08 (m, 1H), 1.82-1.30 (m, 15H), 0.88-0.87 (m, 1H).

1.6.4. Synthesis of 2-amino-6-(((bicyclo[2.2.1]hept-5-en-2-yloxy)carbonyl)amino)hexanoic acid hydrochloride **22**[5]

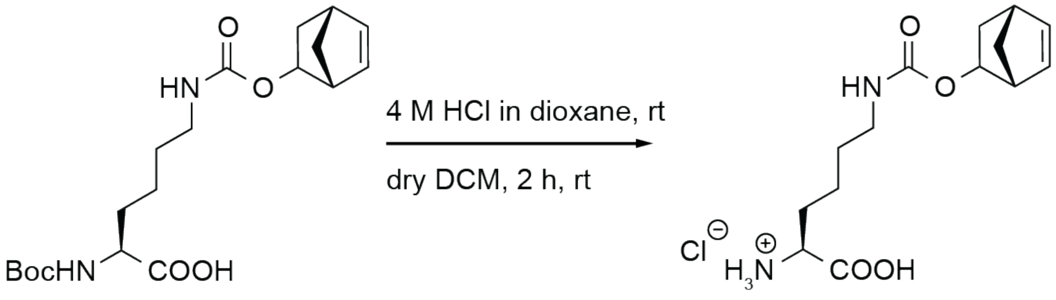


**21** **22**

A solution of the Boc-protected norbornene lysine **21** in DCM (25 mL) and 4 M HCl in dioxane (20 mL, 80 mmol, 8 equiv.) was stirred for 1 h at room temperature. After evaporation of the solvent *in vacuo*, the residue was taken up in diethyl ether and filtrated, to afford **22** as colourless solid (3.10 g, 97%).

$^{1}$H-NMR(400 MHz, DMSO-$d_6$) δ (ppm) = 7.08-6.91 (m, 1H), 6.31-6.24 (m, 1H), 6.00-5.92 (m, 1H), 5.11-5.07 and 4.44-4.42 (m, 1H), 3.86-3.78 (m, 1H), 3.45-3.27 (m, 1H), 3.03 (s, 1H), 2.97-2.88 (m, 2H), 2.81-2.77 (m, 1H), 2.06-2.00 (m, 1H), 1.77-1.75 (m, 2H), 1.59-1.55 (m, 1H), 1.42-1.29 (m, 5H), 0.80-0.77 (m, 1H).

MALDI-MS (m/z): $[M+Na]^+$ calcd for $C_{14}H_{22}N_2O_4$ 305.14718, meas. 305.14741.

#### 1.7. Synthesis of NorLys2 (**26**)

##### 1.7.1. Synthesis of bicyclo[2.2.1]hept-5-en-2-ylmethyl (2,5-dioxopyrrolidin-1-yl) carbonate **24**[9,10]

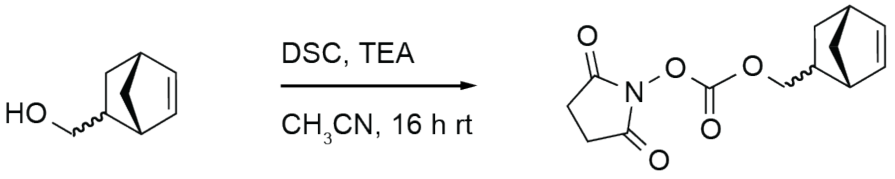


**23** **24**

To a solution of *N,N'*-disuccinimidyl carbonate (13.6 g, 53.6 mmol, 1.1 equiv.) and 5-norbornene-2-methanol **23** (5.8 mL, 48 mmol, 1.0 equiv.) in acetonitrile (120 mL), triethylamine (24 mL) were added, and the mixture was stirred at room temperature overnight. After evaporation of the solvent *in vacuo*, the crude product was purified on silica gel (99.5:0.5. DCM/$Et_2O$) to afford **24** as colourless solid (10.2 g, 80%).

$^1$H-NMR (500 MHz, $CDCl_3$) δ (ppm) = 6.20 (dd, $^3J$ = 3.1 Hz, 5.7 Hz, 1H), 6.13-6.10 (m, 1H), 5.99 (dd, $^3J$ = 2.9 Hz, 5.7 Hz, 1H), 4.41 (dd, $^3J$ = 6.5 Hz, 6.4 Hz, 0.5H), 4.23 (m, 0.5H), 4.11 (dd, $^3J$ = 6.6 Hz, 10.3 Hz, 1H), 3.94 (dd, $^3J$ = 9.8 Hz, 10.1 Hz, 1H), 2.97 (s, 1H), 2.55-2.49 (m, 1H), 1.92-1.83 (m, 2H), 1.50 (dd, $^3J$ = 2.08 Hz, 8.4 Hz, 1H), 1.42-1.18 (m, 3H), 0.61-0.57 (m, 1H).

ESI-MS (m/z): $[M+Na]^+$ calcd for $C_{13}H_{15}NO_5$ 288.25, meas. 288.16.

1.7.2. Synthesis of 6-(((bicyclo[2.2.1]hept-5-en-2-ylmethoxy)carbonyl)amino)-2-((tert-butoxycarbonyl) amino)hexanoic acid **25**[9,10]

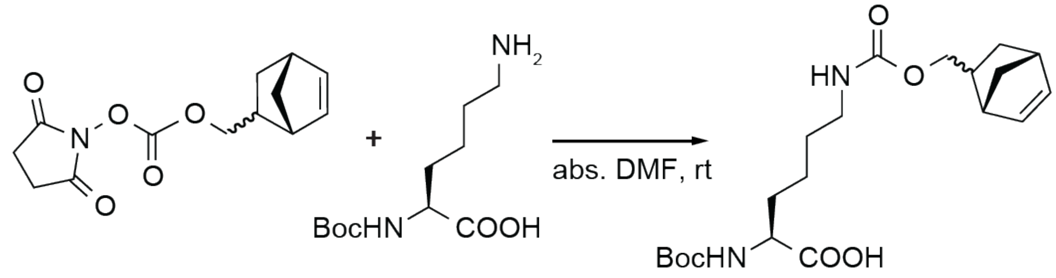


**24** **13** **25**

A solution of compound **24** (9.66 g, 36.4 mmol, 1.0 equiv.) and Boc-protected lysine **13** (9.88 g, 40.0 mmol, 1.1 equiv.) in DMF (100 mL) was stirred at room temperature overnight. After addition of water (400 mL), the mixture was extracted with ethyl acetate (500 mL). The organic phase was washed with water (150 mL) and subsequently, the aqueous phase was extracted twice with ethyl acetate (400 mL). The organic phases were combined and washed again with water (400 mL) and brine (200 mL), and dried over $MgSO_4$. Evaporation of the solvent *in vacuo* afforded **25** as slightly yellow foam (14.4 g, 99%).

$^1$H-NMR (500 MHz, $CDCl_3$) δ (ppm) = 6.17-12 (m, 1H), 6.11-6.06 (m, 1H), 5.95-5.94 (m, 1H), 5.29-5.27 (m, 1H), 4.36-4.26 (m, 1H), 3.20-3.17 (m, 3H), 2.98 (s, 1H), 2.89-2.85 (m, 1H), 1.92-1.80 (m, 3H), 1.57-1.51 (m, 3H), 1.48-1.39 (m, 14H).

ESI-MS (m/z): $[M+H]^+$ calcd for $C_{20}H_{32}N_2O_6$ 397.48, meas. 397.25.

1.7.3. Synthesis of 2-amino-6-(((bicyclo[2.2.1]hept-5-en-2-ylmethoxy)carbonyl)amino)hexanoic acid hydrochloride **26**[5]

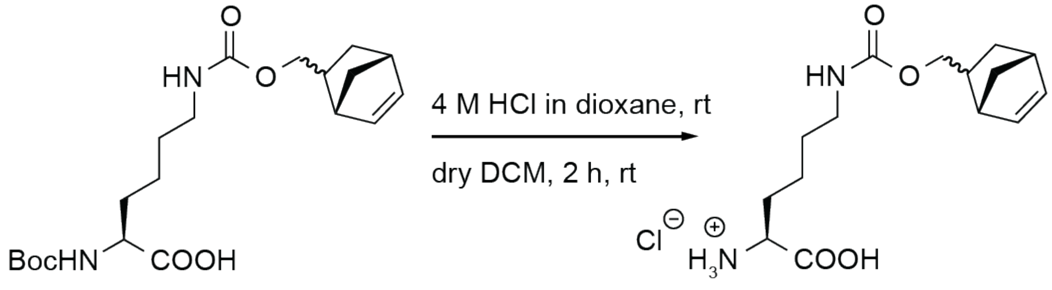


**25** **26**

To a solution of the Boc-protected intermediate **25** (14.4 g, 36.0 mmol, 1.0 equiv.) in DCM (100 mL), 4 M HCl in dioxane (80.0 mL, 0.32 mol, 8.8 equiv.) were added and the solution was stirred for 1 h at room temperature. Evaporation of the solvent *in vacuo* afforded **26** as colourless solid (10.4 g, 87%).

$^1$H-NMR (250 MHz, DSMO-$d_6$) δ (ppm) = 8.31 (m, 2H), 7.14-7.06 (m, 1H), 6.20-6.17 (m, 1H), 6.11-6.09 (m, 1H), 5.95-5.93 (m, 1H), 4.38-4.34 (m, 0.3H), 4.07-4.01 (m, 1H), 2.90 (s, 0.7H), 2.82-2.80 (m, 2H), 2.25-2.23 (m, 1H), 1.86-1.70 (m, 8H), 1.16-1.10 (m, 1H).

MS-ESI (m/z): $[M+H]^+$ calcd for $C_{15}H_{24}N_2O_4$ 297.36, meas. 297.20.

#### 2. Detailed cloning procedure

##### 2.1. Cloning of $pAC^U$ vectors

The $pAC^U$ vector was assembled from cassettes amplified from different commercially available vectors (see Supplementary Fig. S1). The backbone derived from the pCDF-1b vector (EMD Millipore, Novagen), and was cloned as insert with primers CH02 & CH03. The tac promoter and rrnB terminator region derived from the pMAL-c5G vector (NEB, discontinued), and were cloned as inserts with primers CH04 (with 5'-overhang to the pCDF-1b backbone insert) & CH05 and CH32 & CH19.

###### 2.1.1. *Methanococcus jannaschii* derived TyrRS-$tRNA^{Tyr}$ pair

The genes encoding mjTyrRS and $tRNA^{Tyr}$ were cloned as inserts from pEVOL-pBpF with primers CH06 (with 5'-overhang to the tac promoter region from pMAL-c5G) & CH33 (with 5'-overhang to the rrnB terminator region from pMAL-c5G), and CH34 (with 5'-overhang to the rrnB terminator region from pMAL-c5G) & CH11 (with 5'-overhang to the pCDF-1b backbone insert). The vector pEVOL-pBpF was a gift from Peter Schultz (Addgene plasmid # 31190) and contains genes encoding an optimized $tRNA^{Tyr}$ and an evolved mjTyrRS for the incorporation of benzoylphenylalanine. Phusion polymerase (Clontech) was used to generate PCR fragments, which were assembled with help of complementary primer overhang in a MegaPrimer PCR and subsequently cloned into the backbone using InFusion Cloning (Takara Bio). The resulting vector $pAC^U$_Bpa was used as template for multiple point mutations in the gene of *M. jannaschii* derived tyrosyl-tRNA synthetase to generate a variety of $pAC^U$ vectors encoding evolved mjTyrRSs, specific for incorporation of various substituted phenylalanine ncAAs.

###### 2.1.2. *Methanosarcina mazei/ barkeri* derived PylRS-$tRNA^{Pyl}$ pair

In order to incorporate lysine-derivative ncAAs, a *M. mazei* or *M. barkeri* derived pyrrolysyl-tRNA synthetase and corresponding $tRNA^{Pyl}$ is used. Therefore, the genes encoding mjTyrRS and $tRNA^{Tyr}$ in the $pAC^U$_Bpa vector were exchanged by genes encoding mmPylRS/mbPylRS and $tRNA^{Pyl}$. The insert between the genes of PylRS and $tRNA^{Pyl}$ was cloned from $pAC^{U}$ with primers CH32 & AR366 and contained the rrnB terminator and proK promoter sequences. The $pAC^{U}$ backbone was cloned with primers AR369 & CH05.

The inserts encoding mmPylRS and $tRNA^{Pyl}$ were derived from the plasmid pJZ, which was a gift from Nediljko Budisa and contains genes encoding a wild type $tRNA^{Pyl}$ and a N-terminally Strep-tagged mmPylRS evolved for the incorporation of BCN. The mmPylRS insert was cloned without Strep-tag with primers CH53 & AR365 and the $tRNA^{Pyl}$ insert with primers AR367 & AR368.

Phusion polymerase (Clontech) was used to generate the PCR fragments, that were assembled with the help of complementary primer overhangs in MegaPrimer PCRs and subsequently cloned into the backbone using InFusion Cloning (Takara Bio). The resulting vector $pAC^{U}$_BCN was used as template for multiple point mutations in the gene of *M. mazei* derived pyrrolysyl-tRNA synthetase to generate a variety of $pAC^{U}$ vectors encoding evolved mmPylRS, specific for incorporation of various lysine-derivative ncAA.

The wild type $tRNA^{Pyl}$ contains an unfavourable U:G wobble pair in the anticodon stem, which was mutated to a C:G pair to improve suppression efficiency[11]. This mutagenesis was performed with primers CH57 & CH56.

The mbPylRS insert derived from the pAcBac1.tR4-MbPyl vector, which was a gift from Peter Schultz (Addgene plasmid # 50832) and contains genes encoding an undescribed mbPylRS and two copies of synthetic $tRNA^{Pyl}$ derived from *Desulfitobacterium hafniense*. The mbPylRS insert was cloned with primers CH51 & CH52. Same improved $tRNA^{Pyl}$ was used as for the *M. mazei* derived PylRS-$tRNA^{Pyl}$ pair.

Phusion polymerase (Clontech) was used to generate the PCR fragments, that were assembled with the help of complementary primer overhangs in MegaPrimer PCRs and subsequently cloned into the backbone using InFusion Cloning (Takara Bio) resulting in the $pAC^{U}$_PyLys vector. The *M. barkeri* derived pyrrolysyl-tRNA synthetase was only used in its wild type form.

#### 2.2. Cloning of pAC$^{E}$ vectors

Genes encoding different mutants of aaRSs in the pAC$^{U}$ vectors were used as templates to create pAC$^{E}$ plasmids (see Supplementary Fig. S1). The pAC$^{E}$ backbone was derived from the pEVOL-pBpF vector and was amplified with primers CH123 & CH124. The insert between the two copies of the gene encoding aaRS was also derived from the pEVOL-pBpF vector and was cloned with primers CH100 & CH101.

##### 2.2.1. *Methanococcus jannschii* derived TyrRS-tRNA$^{Tyr}$ pair

The two copies of the gene encoding mjTyrRS were derived from the corresponding pAC$^{U}$ plasmids and were amplified with primers CH102 & CH103 and CH104 & CH105, and combined with the pEVOL insert in a MegaPrimer PCR using primers CH116 & CH118, and subsequently cloned into the backbone using InFusion Cloning (Takara Bio).

##### 2.2.2. *Methanosarcina mazei/ barkeri* derived PylRS-tRNA$^{Pyl}$ pair

For the *M. mazei* and *M. barkeri* pAC$^{E}$ plasmids, the gene encoding tRNA$^{Tyr}$ had to be replaced with the respective gene for tRNA$^{Pyl}$. Therefore, primers AR369 & AR366 were used to amplify the pAC$^{E}$ backbone and primers AR367 & AR368 were for the tRNA$^{Pyl}$ insert, respectively. Again, the two copies of the gene encoding both aaRSs were derived from the corresponding pAC$^{U}$ plasmids. For mmPylRS primers CH96 & CH97, and CH98 & CH99 were used to amplify a fragment, which was combined with the pEVOL insert in a MegaPrimer PCR using primers CH115 & CH117, and subsequently cloned into the backbone using InFusion Cloning (Takara Bio).

To amplify the gene encoding the mbPylRS primers CH106 & CH107 and CH108 & CH109 were used and the respective PCR product was combined with the pEVOL insert in a MegaPrimer PCR using primers CH115 & CH119, and subsequently cloned into the backbone using InFusion Cloning (Takara Bio).

#### Supplementary Data:

##### Mass spectrometric analysis of ACP-GFP constructs

Mass spectrometric analysis of purified ACP-GFP constructs was performed on a nanoESI (Synapt G2-S). For each construct the highest peak of the +12 ionized species was picked to calculate the protein mass. Within accuracy of the mass spectrometry device, found protein masses were typically 8-21 Da smaller than theoretical protein masses (except for ACP-GFP_Gln70AzPhe which showed a mass difference of 54 Da). The experimental mass differences between wild type and ACP-GFP with incorporated ncAAs were determined with a deviation of ±7 Da (except for ACP-GFP_Gln70AzPhe which showed a deviation of -39 Da).

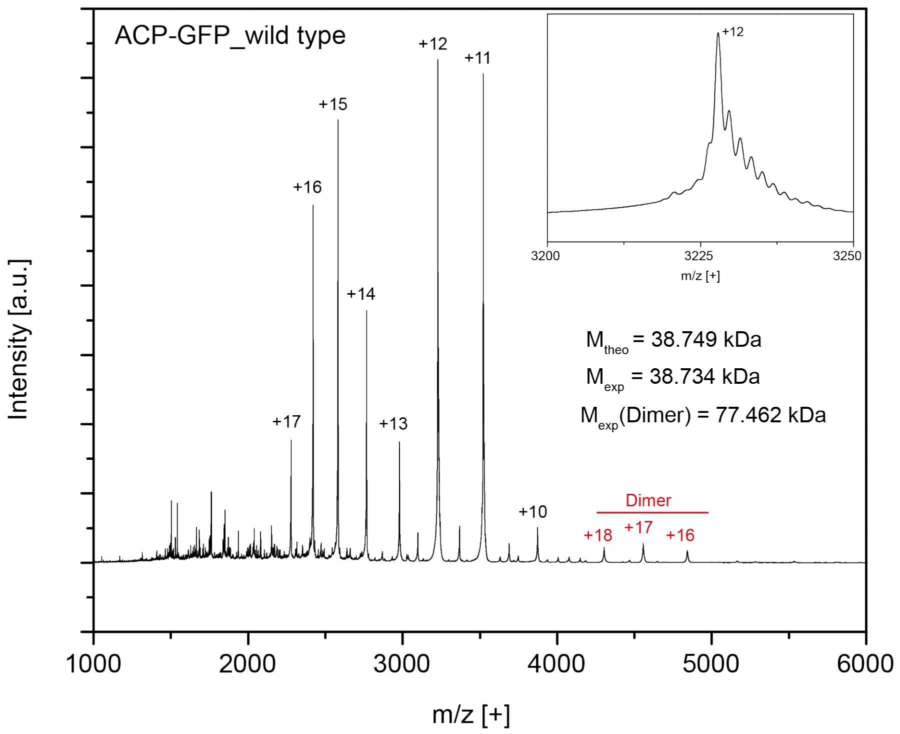
ACP-GFP_wild type
Intensity [a.u.]
m/z [+]
+12
+11
+15
+16
+14
+17
+13
+10
Dimer
+18
+17
+16
$M_{theo}$ = 38.749 kDa
$M_{exp}$ = 38.734 kDa
$M_{exp}$(Dimer) = 77.462 kDa
1000 2000 3000 4000 5000 6000
3200 3225 3250
m/z [+]

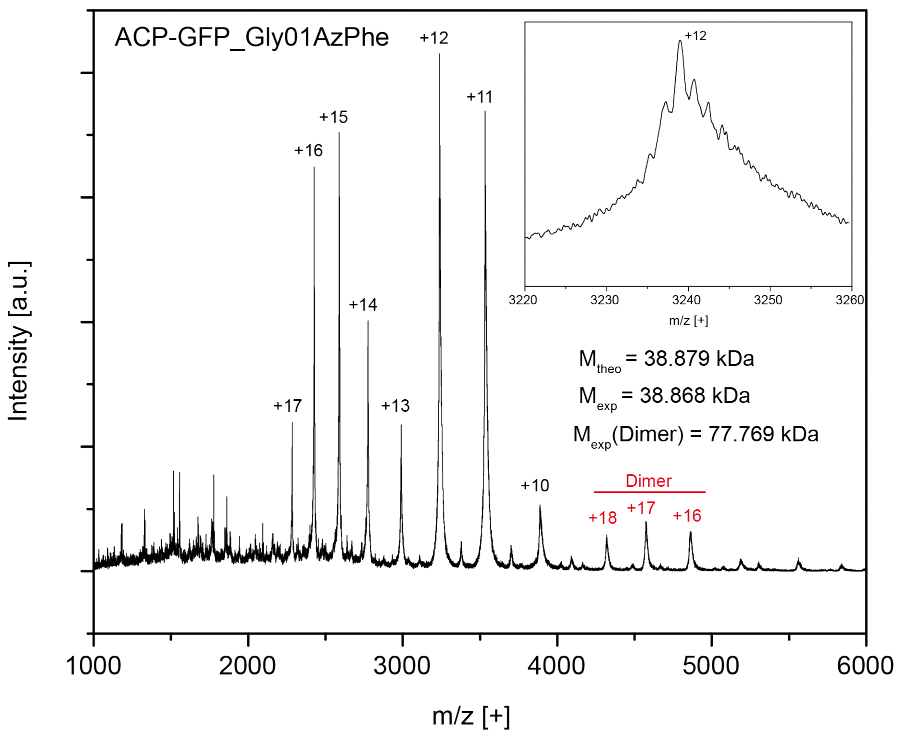
ACP-GFP_Gly01AzPhe
Intensity [a.u.]
m/z [+]
+12
+11
+15
+16
+14
+17
+13
+10
Dimer
+18
+17
+16
$M_{theo}$ = 38.879 kDa
$M_{exp}$ = 38.868 kDa
$M_{exp}$(Dimer) = 77.769 kDa
1000 2000 3000 4000 5000 6000
3220 3230 3240 3250 3260
m/z [+]

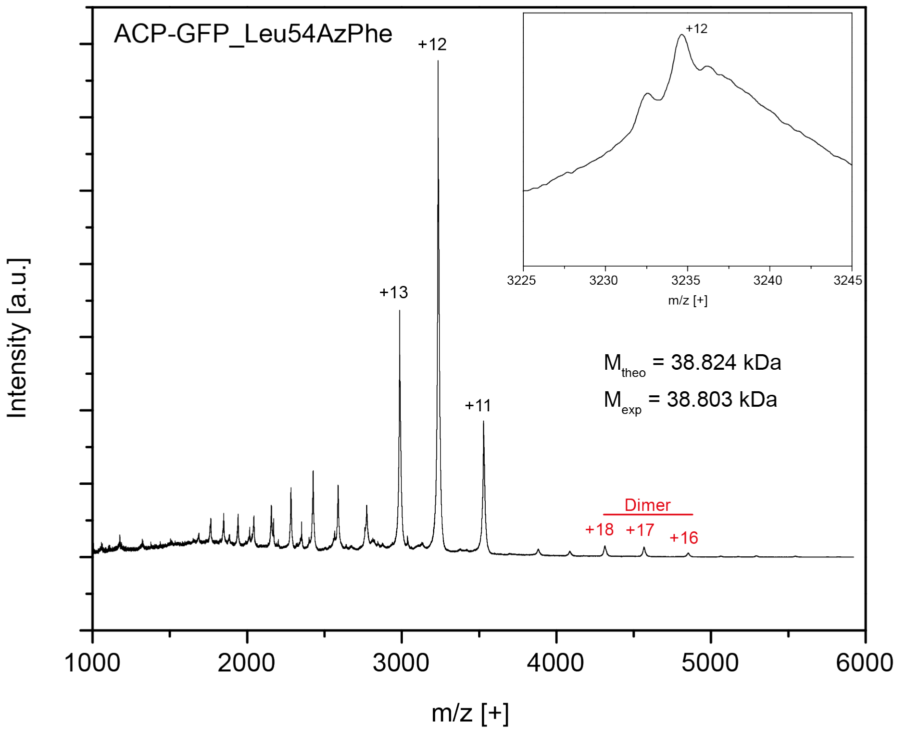
ACP-GFP_Leu54AzPhe
Intensity [a.u.]
m/z [+]
+12
+13
+11
Dimer
+18
+17
+16
M_theo = 38.824 kDa
M_exp = 38.803 kDa
1000
2000
3000
4000
5000
6000
3225
3230
3235
3240
3245
m/z [+]

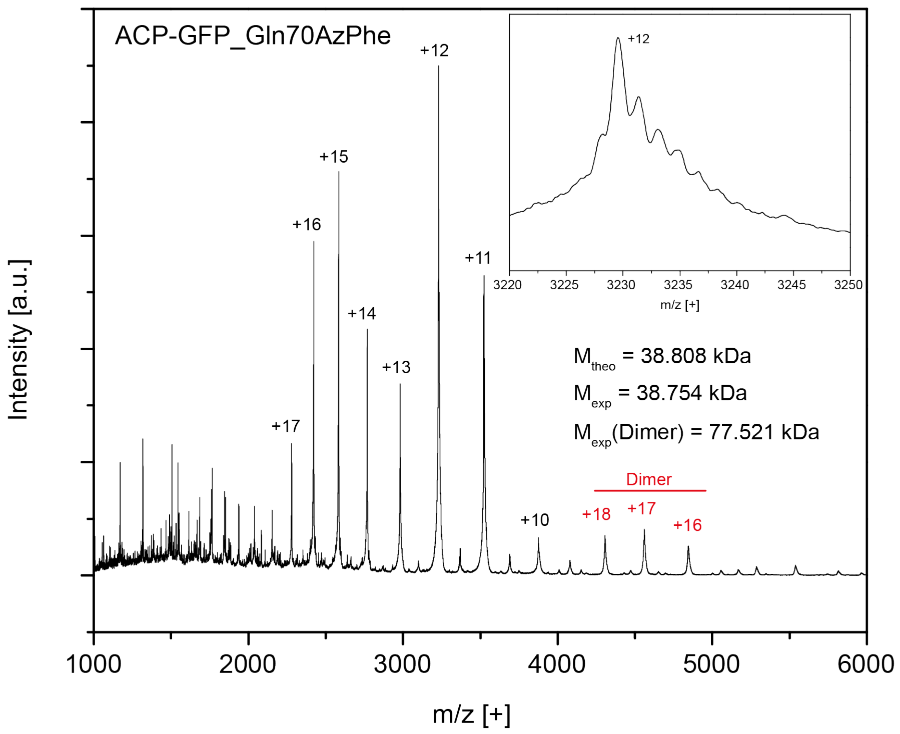
ACP-GFP_Gln70AzPhe
Intensity [a.u.]
m/z [+]
+12
+15
+16
+14
+13
+17
+11
+10
Dimer
+18
+17
+16
M_theo = 38.808 kDa
M_exp = 38.754 kDa
M_exp(Dimer) = 77.521 kDa
1000
2000
3000
4000
5000
6000
3220
3225
3230
3235
3240
3245
3250
m/z [+]

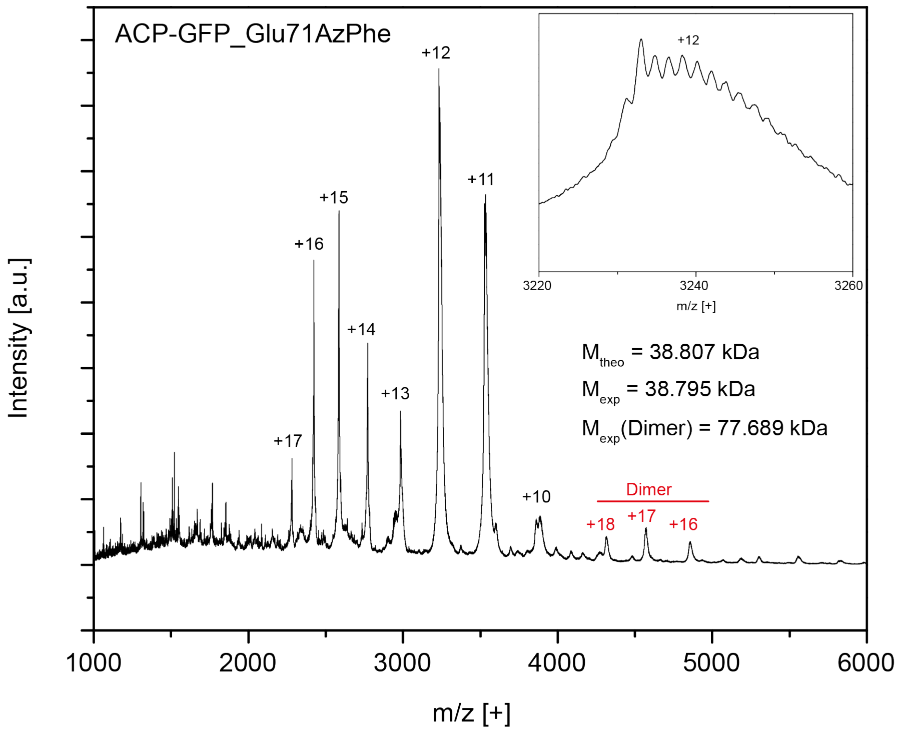
ACP-GFP_Glu71AzPhe
Intensity [a.u.]
m/z [+]
1000
2000
3000
4000
5000
6000
+17
+16
+15
+14
+13
+12
+11
+10
Dimer
+18
+17
+16
3220
3240
3260
m/z [+]
+12
M_theo = 38.807 kDa
M_exp = 38.795 kDa
M_exp(Dimer) = 77.689 kDa

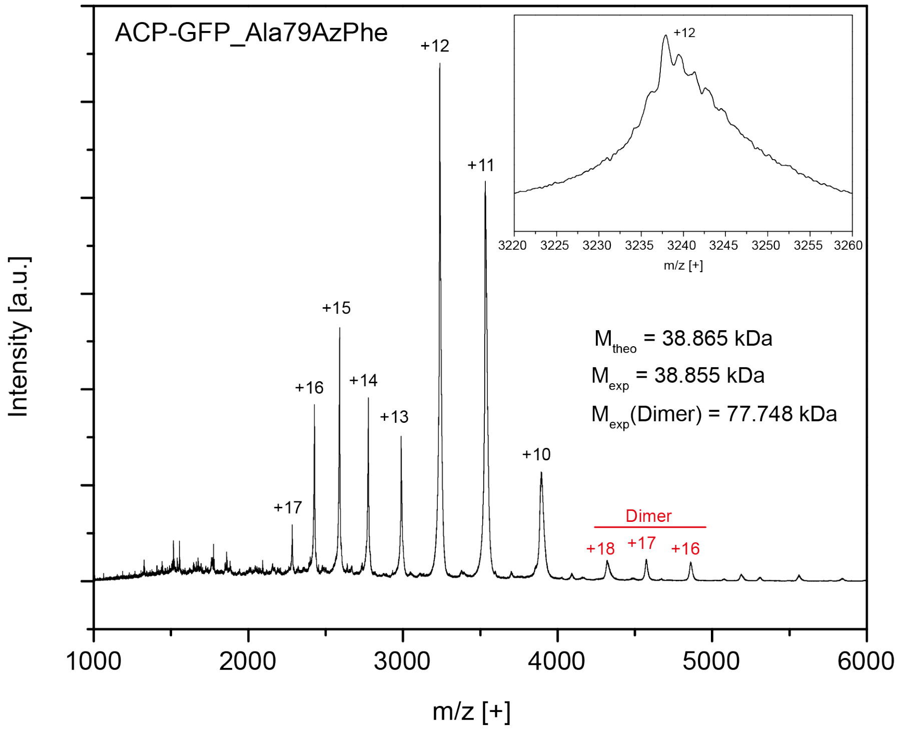
ACP-GFP_Ala79AzPhe
Intensity [a.u.]
m/z [+]
1000
2000
3000
4000
5000
6000
+17
+16
+15
+14
+13
+12
+11
+10
Dimer
+18
+17
+16
3220 3225 3230 3235 3240 3245 3250 3255 3260
m/z [+]
+12
M_theo = 38.865 kDa
M_exp = 38.855 kDa
M_exp(Dimer) = 77.748 kDa

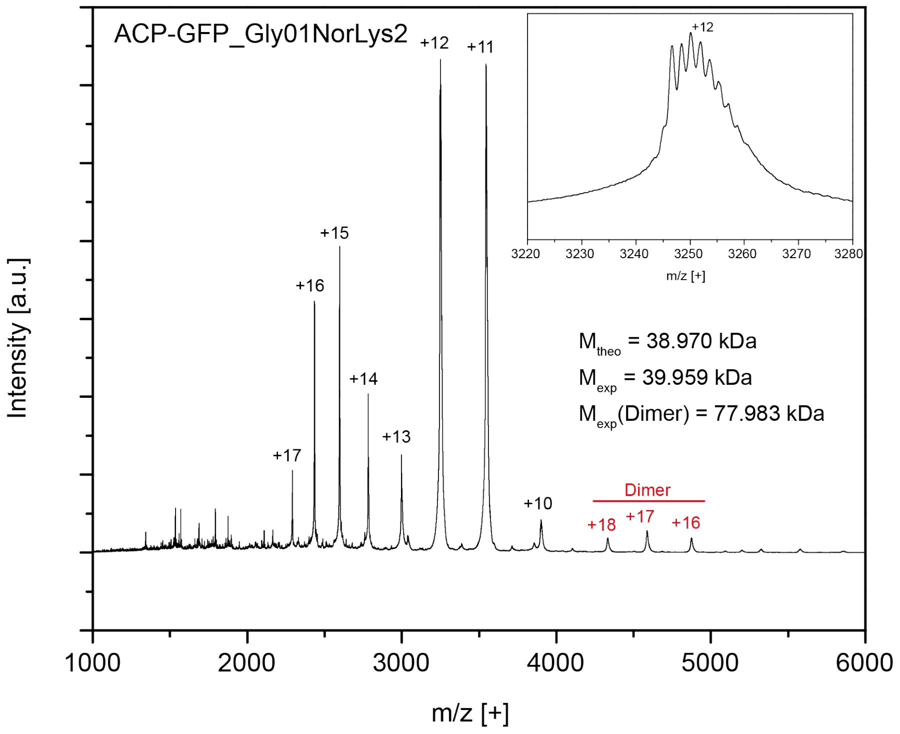

ACP-GFP_Gly01NorLys2
+12
+11
+15
+16
+14
+13
+17
+10
Dimer
+18
+17
+16
Mtheo = 38.970 kDa
Mexp = 39.959 kDa
Mexp(Dimer) = 77.983 kDa
Intensity [a.u.]
m/z [+]
1000
2000
3000
4000
5000
6000


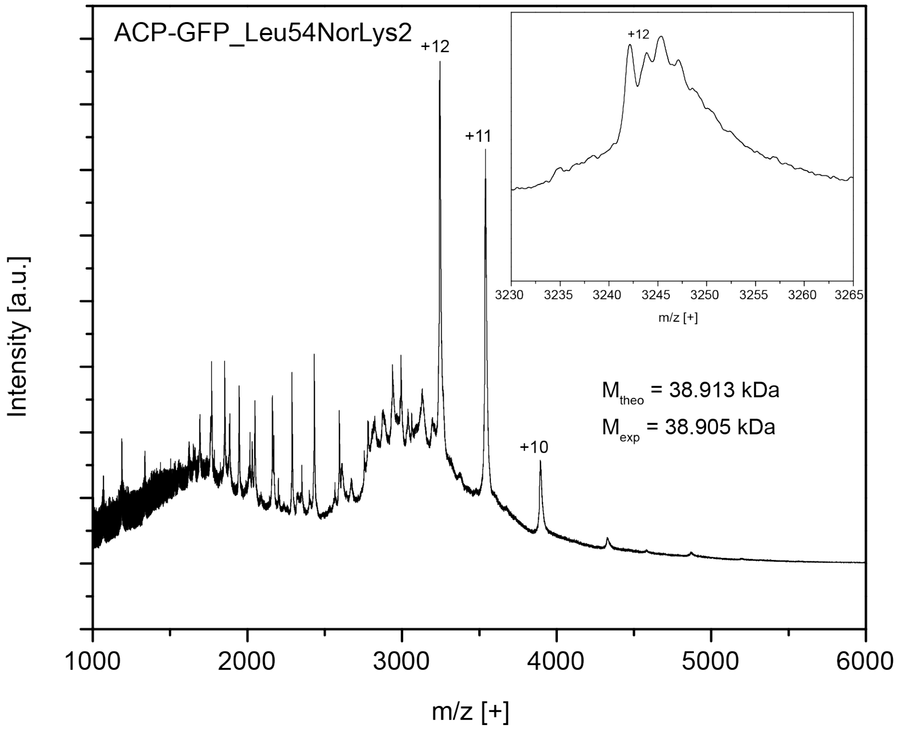

ACP-GFP_Leu54NorLys2
+12
+11
+10
Mtheo = 38.913 kDa
Mexp = 38.905 kDa
Intensity [a.u.]
m/z [+]
1000
2000
3000
4000
5000
6000

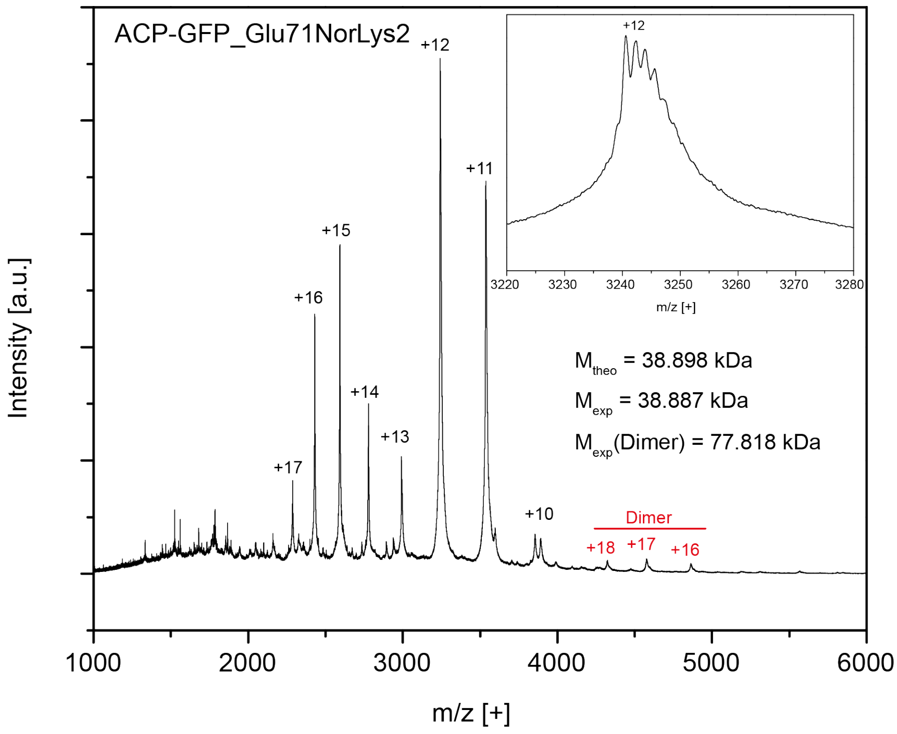
ACP-GFP_Glu71NorLys2
+12
+11
+15
+16
+14
+13
+17
+10
Dimer
+18 +17 +16
$M_{theo}$ = 38.898 kDa
$M_{exp}$ = 38.887 kDa
$M_{exp}$(Dimer) = 77.818 kDa
Intensity [a.u.]
m/z [+]
1000 2000 3000 4000 5000 6000
3220 3230 3240 3250 3260 3270 3280
m/z [+]

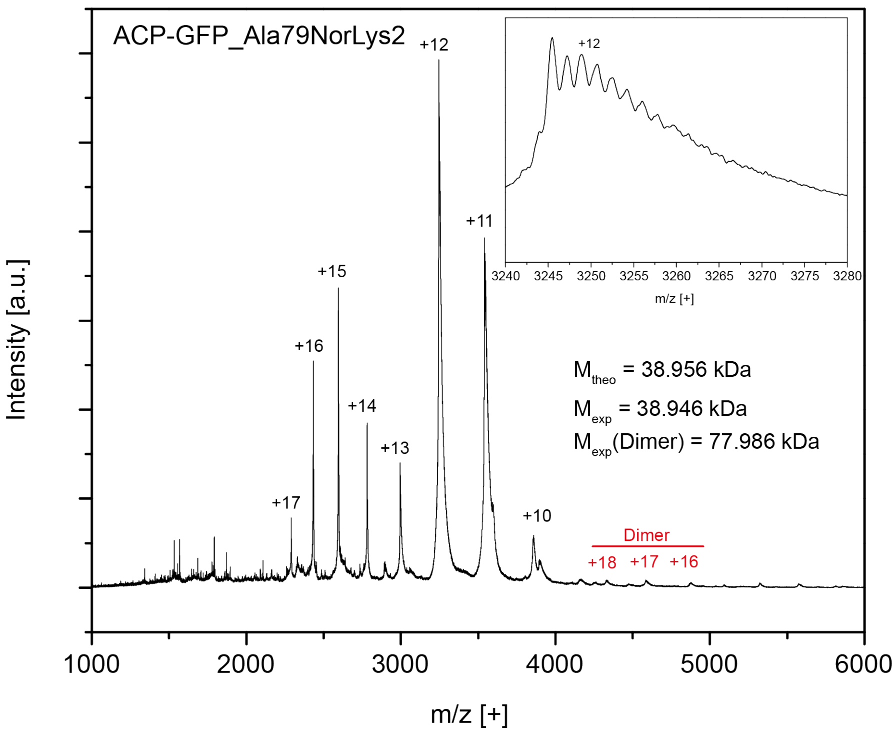
ACP-GFP_Ala79NorLys2
+12
+11
+15
+16
+14
+13
+17
+10
Dimer
+18 +17 +16
$M_{theo}$ = 38.956 kDa
$M_{exp}$ = 38.946 kDa
$M_{exp}$(Dimer) = 77.986 kDa
Intensity [a.u.]
m/z [+]
1000 2000 3000 4000 5000 6000
3240 3245 3250 3255 3260 3265 3270 3275 3280
m/z [+]

#### Supplementary Figures

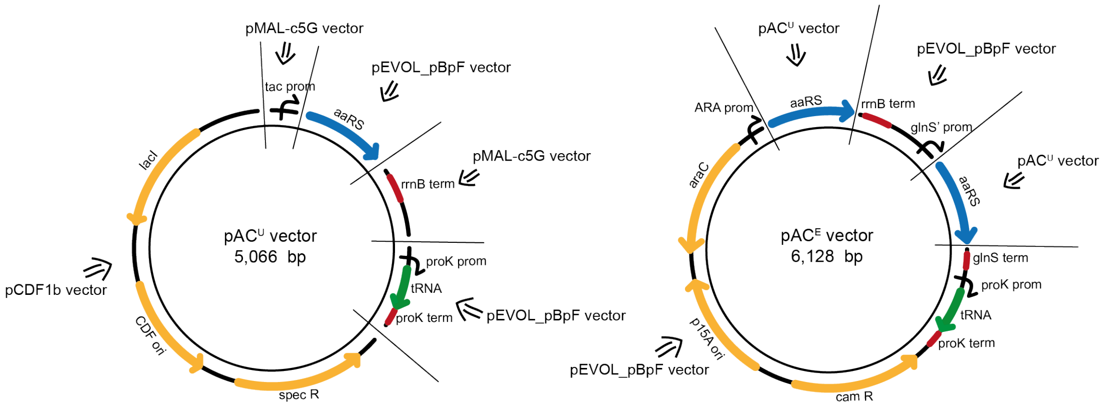


**Figure S1: Cloning scheme of pAC$^{U}$ and pAC$^{E}$ vectors.** The pAC$^{U}$ vectors were constructed using the commercially available vectors pCDF1b, pMAL-c5G and pEVOL_pBpF (for mjTyrRS/tRNA$^{Tyr}$ pair), pJZ (for mmPylRS/tRNA$^{Pyl}$ pair) or pAcBac1.tR4-MbPyl (for mbPylRS/tRNA$^{Pyl}$) as templates. PCR fragments were assembled with help of complementary primer overhangs in MegaPrimer PCRs and subsequently cloned into the backbone using InFusion Cloning (Takara Bio). Multiple point mutations were introduced to create a set of genes encoding the different mutated aaRSs. The amber suppression library of pAC$^{U}$ vectors was used as template to create pAC$^{E}$ plasmids by replacing the genes encoding both copies of the aaRS in pEVOL_pBpF with the genes encoding respective mutants of aaRSs, using the same cloning methods.

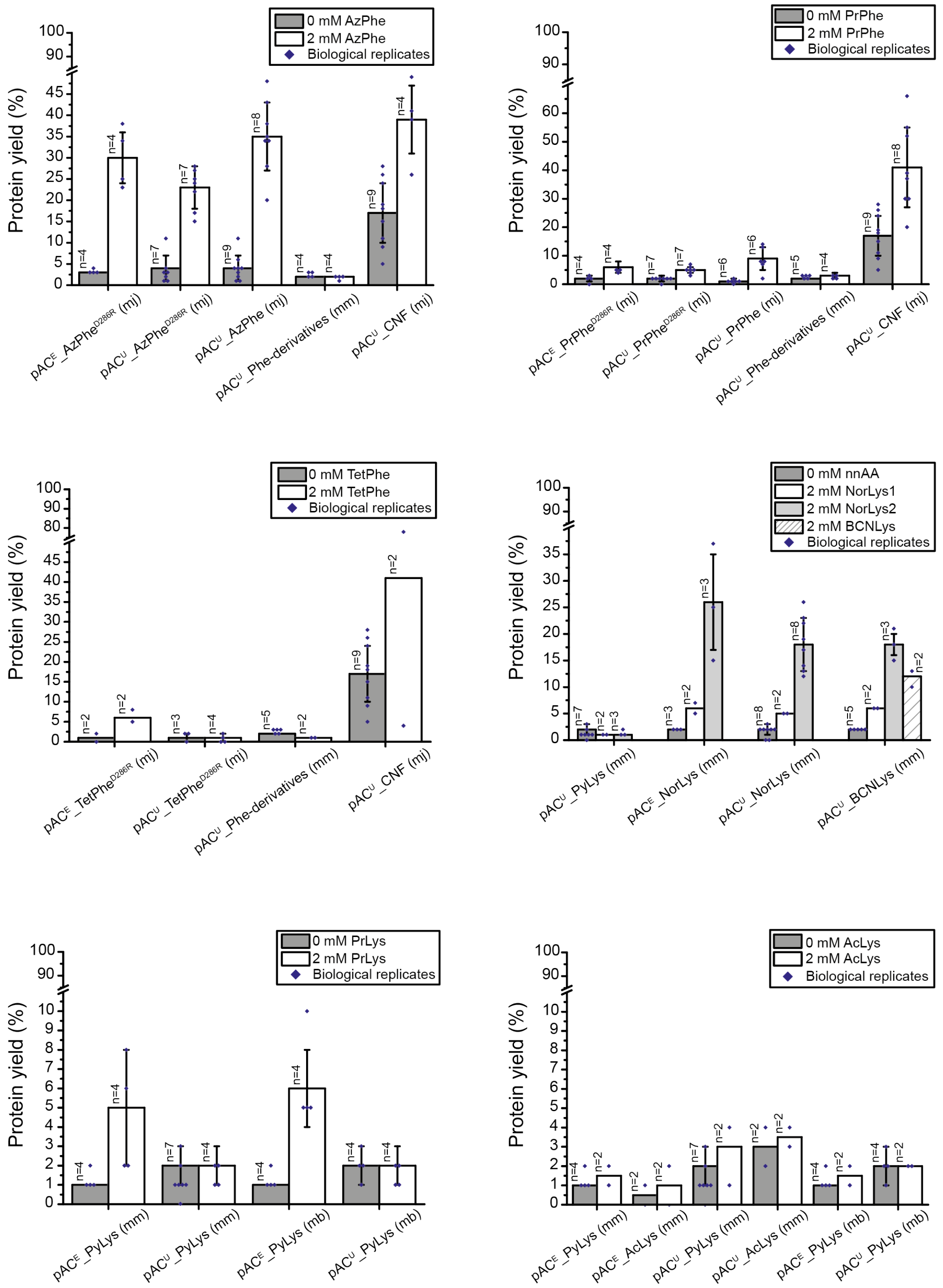


**Figure S2: Screening for the most efficient suppression vector system to introduce select ncAAs with the reporter assay.** The ACP-GFP construct with amber codon at site Leu54 was used for screening. Expression efficiency was read out by GFP fluorescence of 2 mL cell cultures and compared to the wild type reference. For incorporation, 2 mM ncAAs were supplemented to the medium. Cultures lacking ncAAs were taken as negative control to determine background signal. The averages of biological replicates are plotted together with standard deviation and the distribution of individual values is indicated as dots. Technical errors were below 10%.

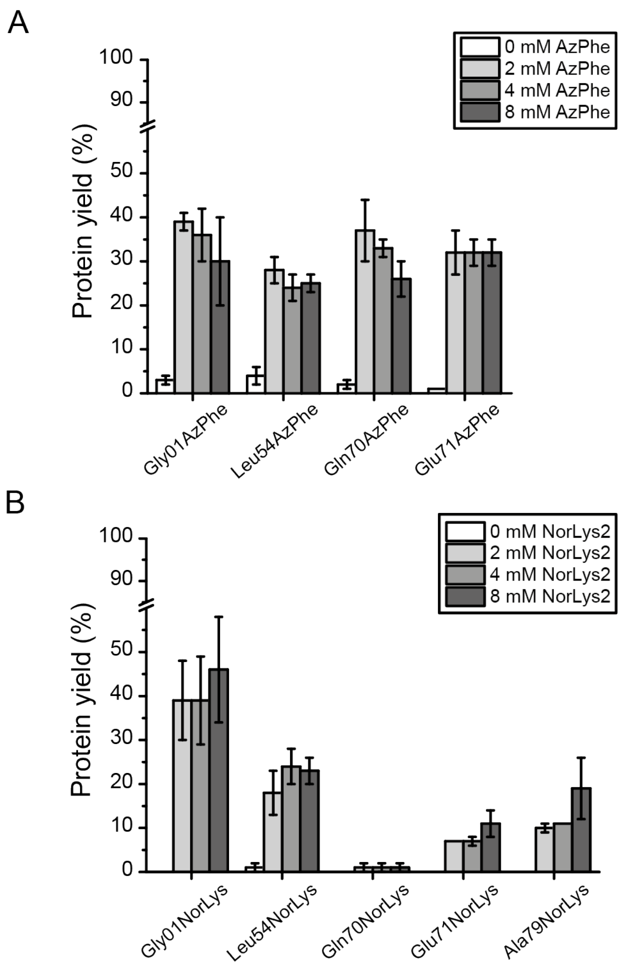


**Figure S3: Testing the effect of different ncAA concentrations.** Incorporation at different sites in the ACP-GFP construct was investigated with the reporter assay in 2 mL scale. Concentrations of 0, 2, 4 and 8 mM ncAA were supplemented to the medium. Expression efficiency was read out by GFP fluorescence and compared to the wild type reference. Error bars reflect the standard deviation of technical triplicates.

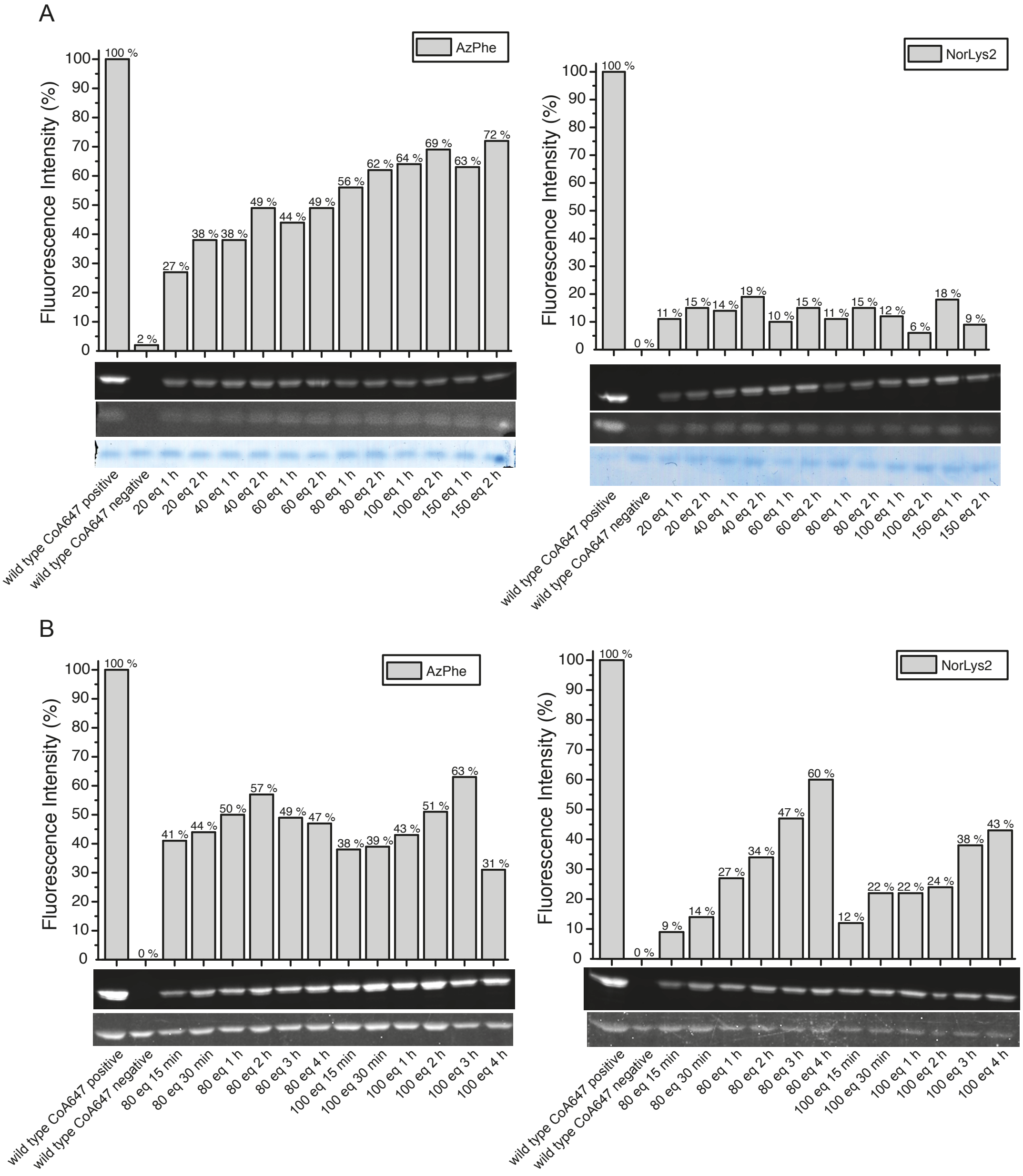


**Figure S4: Optimizing the click reaction conditions for fluorescent labelling of ACP-GFP mutants.** AzPhe mutants were labelled with BCN-POE$_3$-NH-DY649P1, NorLys2 mutants were labelled with 6-methyl-tetrazine-ATTO-647N and the wild type ACP-GFP was modified enzymatically with a fluorescent CoA647-label by Sfp. DOL determined by relative in-gel fluorescence intensities at wavelength 650 nm compared to wild type ACP-GFP reference. All fluorescence intensities were corrected by the quantum efficiencies of the respective fluorophores and correlated to the protein bands of the Syprored- or Coomassie-stained gel. A) DOL of AzPhe mutants (left panel) and NorLys2 mutants (right panel) monitored in dependence of different fluorophore equiv. after labelling reactions for 1 h and 2 h. B) Time dependence of the DOL of AzPhe mutants (left panel) and NorLys2 mutants (right panel) monitored for labelling reactions with 80 and 100 equiv. of fluorophore.

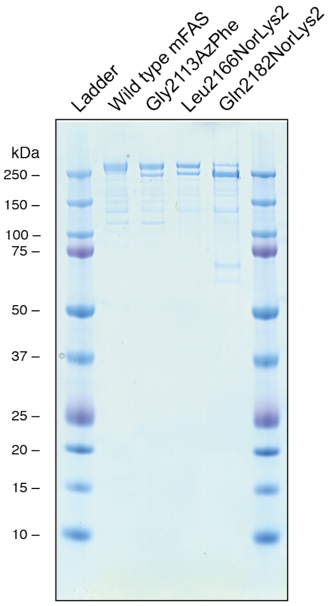


**Figure S5: Generation of ncAA-modified mFAS mutants for fluorescent labelling.** SDS-PAGE (NuPAGE Bis-Tris 4-12%) gel of mFAS mutants. Proteins were purified by a tandem affinity chromatography via the N-terminal Strep-tag and the C-terminal His-tag.

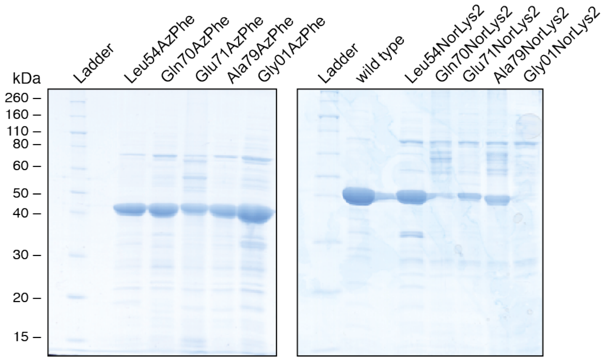


**Figure S6: Large scale expression and purification of ACP-GFP mutants.** Original SDS-PAGE (NuPAGE Bis-Tris 4-12%) gel of ACP-GFP mutants purified by Ni-chelating chromatography. Referring to Fig. 4 in the manuscript.

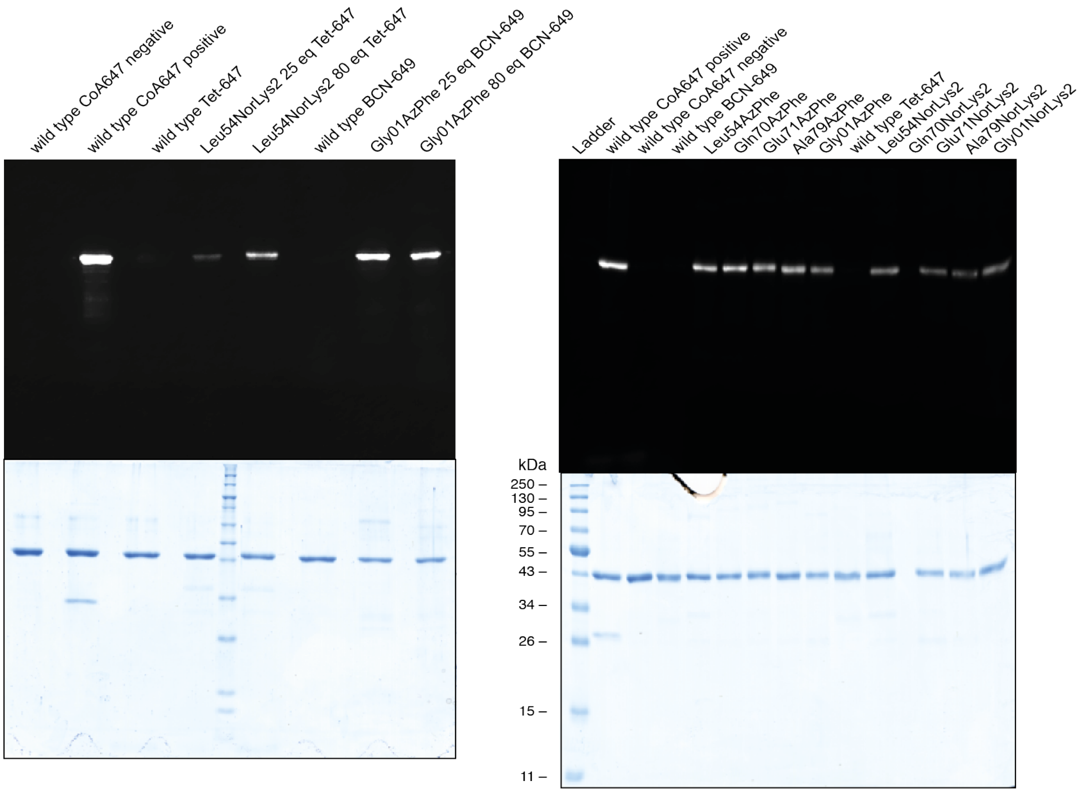


**Figure S7: Fluorescent labelling of ACP-GFP mutants.** Original fluorescent gel and Coomassie-stained gel of fluorescently labelled ACP-GFP mutants. Referring to Fig. 5 in the manuscript.

#### Supplementary Tables

Table S1: List of primers used for cloning amber codon suppression vectors $pAC^U$ and $pAC^E$.

| # | Primer name | Primer sequence |
|---|---|---|
| CH02 | pCDF1b_for | CTGAAACCTCAGGCATTTGAGAAG |
| CH03 | pCDF1b_rev | ATTTCCTAATGCAGGAGTCGCATAAG |
| CH04 | pCDF_tac_for | CCTGCATTAGGAAATTTGACAATTAATCATCGGCTCGTATAATG |
| CH05 | tac_Nde1-rev | ATGCTATGGTCCTTGTTGGTCAATTGC |
| CH06 | tac_mjTyrS_for | CAATTGACCAACAAGGACCATAGCATATGGACGAATTTGAAATGATAAAGAGAAAC |
| CH11 | mjtRNA_pCDF1b_rev | TGCCTGAGGTTTCAGCAAATTCGACCCTGAGCTGC |
| CH19 | mjtRNA_rrnBterm_rev | GCGGAATTATTTGATCGGCGAGGTCTCATGAGCGGATACATATTTG |
| CH32 | rrnBterm_for | GGTGCGGCACACTACGTAC |
| CH33 | mjTyrS_rrnBterm_rev | GTACGTAGTGTGCCGCACCTTATAATCTCTTTCTAATTGGCTCTAAAATC |
| CH34 | rrnBterm_mjtRNA_for | CTCGCCGATCAAATAATTCCGCTAATTCCGCTTCGCAACATGTGAG |
| CH51 | tac_mbPylRS | CAAGGACCATAGCATATGGATAAAAAACCATTAGATGTTTTAATATCTGC |
| CH52 | mbPylRS_rrnBterm_rev | GTAGTGTGCCGCACCTCATAGATTGGTTGAAATCCCATTATAGTAAG |
| CH53 | tac_notag-mmPylRS_for | CAAGGACCATAGCATATGGATAAAAAACCACTAAACACTCTGATATC |
| CH56 | PyltRNA_wobble_rev | TTCGATCTACATGATCAGGTTTCCAATG |
| CH57 | PyltRNA_wobble_for | ATCATGTAGATCGAAcGGACTCTAAATCCGTTCAGCC |
| CH96 | ara_mmPylRS_forward | GGAGGAATTAGATCTATGGATAAAAAACCACTAAACACTCTGATATCTG |
| CH97 | mmPylRS_rrnB_rev | CTCAATGATGATGATGATGATGGTCGACTTATTACAGGTTGGTAGAAATCCCGTTATAG |
| CH98 | glnsprom_mmPylRS_for | CGTTGTTTACGCTTTGAGGAATCCCATATGGATAAAAAACCACTAAACACTCTGATATCTG |
| CH99 | mmPylRS_glnsterm_reverse | CGTTTGAAACTGCAGTTACAGGTTGGTAGAAATCCCGTTATAG |
| CH100 | pEVOL_insert_forward | TAAGTCGACCATCATCATCATCATC |
| CH101 | pEVOL_insert_reverse | ATGGGATTCCTCAAAGCGTAAACAAC |
| CH102 | ara_mjTyrRS_forward | GGAGGAATTAGATCTATGGACGAATTTGAAATGATAAAGAGAAAC |
| CH103 | mjTyrRS_rrnB_rev | CTCAATGATGATGATGATGATGGTCGACttaTTATAATCTCTTTCTAATTGGCTCTAAAATCTTTA |
| CH104 | glnsprom_mjTyrRS_for | CGTTGTTTACGCTTTGAGGAATCCCATATGGACGAATTTGAAATGATAAAGAGAAAC |
| CH105 | mjTyrRS_glnsterm_rev | CGTTTGAAACTGCAGTTATAATCTCTTTCTAATTGGCTCTAAAATCTTTA |
| CH106 | ara_mbPylRS_for | GGAGGAATTAGATCTATGGATAAAAAACCATTAGATGTTTTAATATCTG |
| CH107 | mbPylRS_rrnBterm_rev | CTCAATGATGATGATGATGATGGTCGACTTATCATAGATTGG |

| | | |
|---|---|---|
| | | TTGAAATCCCATTATAG |
| CH108 | glnsprom_mbPylRS_for | CGTTGTTTACGCTTTGAGGAATCCCATATGGATAAAAAACCATTAGATGTTTTAATATCTG |
| CH109 | mbPylRS_glnsterm_rev | CGTTTGAAACTGCAGTCATAGATTGGTTGAAATCCCATTATAG |
| CH115 | MegaPrimer_mmpEvol_for | GGAGGAATTAGATCTATGGATAAAAAAC |
| CH116 | MegaPrimer_mjpEvol_for | GGAGGAATTAGATCTATGGACGAATTTG |
| CH117 | MegaPrimer_mmpEvol_rev | CGTTTGAAACTGCAGTTACAGGTTG |
| CH118 | MegaPrimer_mjpEvol_rev | CGTTTGAAACTGCAGTTATAATCTCTTTC |
| CH119 | MegaPrimer_mbpEvol_rev | CGTTTGAAACTGCAGTCATAGATTGG |
| CH123 | pEvol_backbone_for | CTGCAGTTTCAAACGCTAAATTGCC |
| CH124 | pEvol_backbone_rev | AGATCTAATTCCTCCTGTTAGCCC |
| AR365 | mmPylS_rrnB_rev | GTACGTAGTGTGCCGCACCTTACAGGTTGGTAGAAATCCCGTTATAG |
| AR366 | proK_rev | AATGCGGGGCGCATCTTACTG |
| AR367 | proK_mmtRNA_for | CAGTAAGATGCGCCCCGCATTGGAAACCTGATCATGTAGATCGAATG |
| AR368 | mmtRNA_terK_rev | GCTTTTCGAATTTGGTGGCGGAAACCCCGGGAATC |
| AR369 | terK_for | CCAAATTCGAAAAGCCTGCTCAAC |

Table S2: List of amber codon suppressor plasmids used in this study.

| # | Plasmid name | aaRS variant | ncAA |
|---|---|---|---|
| 1 | pAC$^{U}$_AzPhe (mj) | *Methanococcus jannaschii* tyrosyl-tRNA synthetase[12] Y32T, E107N, D158P, I159L, L162Q | AzPhe |
| 2 | pAC$^{U}$_AzPhe$^{D286R}$ (mj) | *Methanococcus jannaschii* tyrosyl-tRNA synthetase[12,13] Y32T, E107N, D158P, I159L, L162Q, D286R | |
| 3 | pAC$^{E}$_AzPhe$^{D286R}$ (mj) | | |
| 4 | pAC$^{U}$_PrPhe (mj) | *Methanococcus jannaschii* tyrosyl-tRNA synthetase[14] Y32A, E107P, L110F, D158A, L162A | PrPhe |
| 5 | pAC$^{U}$_PrPhe$^{D286R}$ (mj) | *Methanococcus jannaschii* tyrosyl-tRNA synthetase[13,14] Y32A, E107P, L110F, D158A, L162A, D286R | |
| 6 | pAC$^{E}$_PrPhe$^{D286R}$ (mj) | | |
| 7 | pAC$^{U}$_TetPhe$^{D286R}$ (mj) | *Methanococcus jannaschii* tyrosyl-tRNA synthetase[5,13] Y32E, L65A, F108P, Q109S, D158G, L162G, D286R | TetPhe |
| 8 | pAC$^{E}$_TetPhe$^{D286R}$ (mj) | | |
| 9 | pAC$^{U}$_CNF (mj) | *Methanococcus jannaschii* tyrosyl-tRNA synthetase[13,15] Y32L, F108W, Q109M, D158G, I159A, D286R | AzPhe; PrPhe; TetPhe |
| 10 | pAC$^{U}$_PyLys (mm) | *Methanosarcina mazei* pyrrolysyl-tRNA synthetase[16] | NorLys1; NorLys2; PrLys; AcLys |
| 11 | pAC$^{E}$_PyLys (mm) | | |
| 12 | pAC$^{U}$_AcLys (mm) | *Methanosarcina mazei* pyrrolysyl-tRNA synthetase[17] L301M, Y306L, L309A, C348F | AcLys |
| 13 | pAC$^{E}$_AcLys (mm) | | |
| 14 | pAC$^{U}$_NorLys (mm) | *Methanosarcina mazei* pyrrolysyl-tRNA synthetase[18] Y306G, Y384F, I405R | NorLys1; NorLys2 |
| 15 | pAC$^{E}$_NorLys (mm) | | |
| 16 | pAC$^{U}$_BCN (mm) | *Methanosarcina mazei* pyrrolysyl-tRNA synthetase[19] Y306A, Y348A | BCNLys; NorLys1; NorLys2 |
| 17 | pAC$^{U}$_Phe-derivatives (mm) | *Methanosarcina mazei* pyrrolysyl-tRNA synthetase[20] N346A, C348A | AzPhe; PrPhe; TetPhe |
| 18 | pAC$^{U}$_PyLys (mb) | *Methanosarcina barkeri* pyrrolysyl-tRNA synthetase[21] | PrLys; AcLys |
| 19 | pAC$^{E}$_PyLys (mb) | | |

#### References


1 Wieland, T., von Dungen, A. & Birr, C. Synthese einer antitoxischen Antamid-Variante mit *p*-Azido-phenylalanin in Stelle 6. *Liebigs Ann. Chem.* **752**, 109-114 (1971).

2 Lieber, E., Levering, D. R. & Patterson, L. Infrared Absorption Spectra of Compounds of High Nitrogen Content. *Anal. Chem.* **23**, 1594-1604 (1951).

3 Jain, S. & Reiser, O. Immobilization of Cobalt(II) Schiff Base Complexes on Polystyrene Resin and a Study of Their Catalytic Activity for the Aerobic Oxidation of Alcohols. *ChemSusChem* **1**, 534-541, doi:10.1002/cssc.200800025 (2008).

4 Borzilleri, R., Zhang, Y., Miller, M. M. & Seigal, B. A. Preparation of macrocyclic compounds for inhibition of inhibitors of apoptosis. (2014).

5 Seitchik, J. L. *et al.* Genetically Encoded Tetrazine Amino Acid Directs Rapid Site-Specific *in Vivo* Bioorthogonal Ligation with *trans*-Cyclooctenes. *J. Am. Chem. Soc.* **134**, 2898-2901, doi:10.1021/ja2109745 (2012).

6 Fields, S. C., Parker, M. H. & Erickson, W. R. A simple route to unsymmetrically substituted 1, 2, 4, 5-tetrazines. *J. Org. Chem.* **59**, 8284-8287 (1994).

7 Nguyen, D. P. *et al.* Genetic Encoding and Labeling of Aliphatic Azides and Alkynes in Recombinant Proteins *via* a Pyrrolysyl-tRNA Synthetase/$tRNA_{CUA}$ Pair and Click Chemistry. *J. Am. Chem. Soc.* **131**, 8720-8721 (2009).

8 Khan, F. A. & Choudhury, S. An Efficient Synthesis of Substituted *meta*-Halophenols and Their Methyl Ethers: Insight into the Reaction Mechanism. *Eur. J. Org. Chem.* **2010**, 2954-2970 (2010).

9 Lang, K. *et al.* Genetically encoded norbornene directs site-specific cellular protein labelling via a rapid bioorthogonal reaction. *Nat. Chem.* **4**, 298-304, doi:10.1038/nchem.1250 (2012).

10 Kaya, E. *et al.* A Genetically Encoded Norbornene Amino Acid for the Mild and Selective Modification of Proteins in a Copper-Free Click Reaction. *Angew. Chem. Int. Ed.* **51**, 4466-4469, doi:10.1002/anie.201109252 (2012).

11 Chatterjee, A., Sun, S. B., Furman, J. L., Xiao, H. & Schultz, P. G. A versatile platform for single- and multiple-unnatural amino acid mutagenesis in *Escherichia coli*. *Biochemistry* **52**, 1828-1837, doi:10.1021/bi4000244 (2013).

12 Chin, J. W. *et al.* Addition of *p*-Azido-L-phenylalanine to the Genetic Code of *Escherichia coli*. *J. Am. Chem. Soc.* **124**, 9026-9027, doi:10.1021/ja027007w (2002).

13 Kobayashi, T. *et al.* Structural basis for orthogonal tRNA specificities of tyrosyl-tRNA synthetases for genetic code expansion. *Nat. Struct. Biol.* **10**, 425-432 (2003).

14 Deiters, A. & Schultz, P. G. *In vivo* incorporation of an alkyne into proteins in *Escherichia coli*. *Bioorg. Med. Chem. Lett.* **15**, 1521-1524, doi:10.1016/j.bmcl.2004.12.065 (2005).

15 Young, D. D. *et al.* An Evolved Aminoacyl-tRNA Synthetase with Atypical Polysubstrate Specificity. *Biochemistry* **50**, 1894-1900, doi:10.1021/bi101929e (2011).

16 Mukai, T. *et al.* Adding L-lysine derivatives to the genetic code of mammalian cells with engineered pyrrolysyl-tRNA synthetases. *Biochem. Biophys. Res. Commun.* **371**, 818-822 (2008).

17 Umehara, T. *et al.* *N*-Acetyl lysyl-tRNA synthetases evolved by a CcdB-based selection possess *N*-acetyl lysine specificity *in vitro* and *in vivo*. *FEBS Lett.* **586**, 729-733, doi:10.1016/j.febslet.2012.01.029 (2012).

18 Schneider, S. *et al.* Structural Insights into Incorporation of Norbornene Amino Acids for Click Modification of Proteins. *ChemBioChem* **14**, 2114-2118, doi:10.1002/cbic.201300435 (2013).

19 Plass, T. *et al.* Amino Acids for Diels-Alder Reactions in Living Cells. *Angew. Chem. Int. Ed.* **51**, 4166-4170, doi:10.1002/anie.201108231 (2012).

20 Wang, Y.-S., Fang, X., Wallace, A. L., Wu, B. & Liu, W. R. A Rationally Designed Pyrrolysyl-tRNA Synthetase Mutant with a Broad Substrate Spectrum. *J. Am. Chem. Soc.* **134**, 2950-2953, doi:10.1021/ja211972x (2012).

21 Blight, S. K. *et al.* Direct charging of $tRNA_{CUA}$ with pyrrolysine *in vitro* and *in vivo*. *Nature* **431**, 333-335 (2004).
